# An eicosanoid-enriched follicular microenvironment shapes germinal centre immune dynamics in antiretroviral-treated PLWH

**DOI:** 10.64898/2026.09.09.750310

**Authors:** Spiros Georgakis, Helen Lindsay, Michail Orfanakis, Jan Lukas Rinker, Perla Mariana Del Rio Estrada, Riddhima Banga, Cloe Brenna, Senbai Kang, Fernanda Torres-Ruiz, Yara Andrea Luna Villalobos, Gonzalo Salgado Montes de Oca, Matthieu Perreau, Mareva Delporte, Olivier Lambotte, Denarda Dangaj-Laniti, Laurence de Leval, Linos Vandekerckhove, Susan Pereira Ribeiro, Raphael Gottardo, Constantinos Petrovas

## Abstract

Delineating the follicular (F) cellular and molecular landscape in HIV infection is essential for understanding neutralizing antibody responses and HIV reservoir maintenance. Multiplex imaging analysis revealed a less differentiated profile for follicular helper CD4 T cells (T_FH_) and significantly increased follicular CD25^lo/hi^Foxp3^hi^ T cells in lymph nodes (LNs) from antiretroviral treated (cART) compared to viremic (Vir) people living with HIV (PLWH). Spatial transcriptomics identified distinct inflammatory follicular microenvironments in Vir and cART LNs, characterized by interferon and eicosanoid enrichment, respectively. In an independent cohort, scRNA-sequencing analysis of LN-derived cells revealed an enrichment of eicosanoid-related pathways in follicular immune cell types in non-neutralizers compared to neutralizers PLWH. *In vitro* infection and *in situ* multimodal investigation of HIV DNA^+^ cell microenvironments suggested a potential role of PGE_2_/Eicosanoids in maintaining viral reservoirs in cART LNs. Our data highlight cellular and molecular factors that could regulate both antibody responses and viral reservoir persistence in PLWH.

Keywords: Follicular Helper CD4 cells, HIV, germinal centre, interferons, eicosanoids.

## Introduction

Even though combined antiretroviral therapy (cART) leads to undetectable HIV viral loads in people living with HIV (PLWH), the virus cannot be eradicated due to the establishment of viral reservoirs and the impaired anti-HIV responses after ART interruption (1–4). Additionally, vaccine responses are weaker and decline faster in PLWH, even those on cART (5). Therefore, a comprehensive understanding of follicular/germinal centre (F/GC) microenvironment in HIV is of great importance for discovering molecular targets that could modulate these biological processes.

Lymph nodes (LNs) are of particular interest as the primary site for the development of anti-HIV adaptive immune responses (6, 7). Follicles/Germinal centers (F/GCs) are specialized niches where B cells undergo somatic hypermutation and affinity maturation to generate memory B cells and long-lived plasma cells producing high-affinity antibodies (8, 9). T_FH_ cells, a phenotypically and functionally heterogeneous cell subset (10–14), mediate this process which is crucial for producing broadly neutralizing antibodies (bnAbs) (15–19). Upon antiretroviral treatment, HIV uses follicular and extrafollicular LN areas as hideouts to survive in a latent state (20–22). During active HIV and SIV infection, GC-B and T_FH_ cells become hyperactivated, driving profound changes in humoral antiviral responses (16, 23–25). Following ART, active HIV transcription is blocked while low-grade HIV-related inflammation still occurs (26–28). During that stage, T_FH_ cells were reported to be important latent-HIV reservoirs together with other LN cell types (29–31).

CD8 T cells are crucial for eliminating HIV-infected cells (32–35). The exhaustion and the limited access of effector CD8 to F/GCs are linked to uncontrolled HIV inflammation (34, 36–38). During cART, LN CD8 T cells acquire stem-like features associated with enhanced proliferative capacity upon secondary antigen exposure and improved anti-HIV immune responses (39–41). However, CD8 cells do not sufficiently block HIV and SIV rebound upon cART interruption (42–44). LN T regulatory (T_regs_) and T follicular regulatory cells (T_FRs_) can limit deleterious hyper-reactive immune reactions in LNs and fine-tune T_FH_-B cell interactions during untreated-HIV disease but also constrain beneficial anti-HIV responses in both untreated and treated patients (45–47). Innate immune cells in LNs are also crucial regulators of adaptive anti-HIV responses through immunomodulators’ secretion (e.g cytokines and lipid mediators like type I IFNs and eicosanoids) and cell-to-cell interactions (48–52).

Type I and II IFNs are essential for potent anti-viral immune responses early during the infection and their multifaceted role on anti-HIV responses has been thoroughly investigated (53–55). In cART-PLWH and animal models, a transcriptomic IFN signature is accompanied by T-cell exhaustion and viral persistence while IFN-α blockade has been shown to be beneficial in SIV (56–58). On the other hand, non-cytokine regulators (e.g lipids, metabolites) can reshape the LN and follicular milieu during adaptive immune responses (59–64). The role of lipid mediators has not been thoroughly investigated in LN-localized anti-HIV responses. Small-molecule derivatives of arachidonic acid, such as Prostaglandin-E (PGE ), are potent immunomodulators (62, 65). PGE is produced primarily by innate and stromal cells and acts on G protein-coupled E receptors (EP1-4) to mediate context-dependent pro-or anti-inflammatory responses (61, 65, 66). PGE suppresses i) cytotoxic T lymphocytes’ (CTL) responses during chronic LCMV infection, ii) type I IFN production by macrophages during influenza A infection and iii) *in-vitro* HIV replication, infectivity and release of virions post infection (67–70). Several studies have investigated the effect of PGE_2_ signalling blockade in chronic viral infections, including HIV, showing variable outcomes (61, 67, 71–73). Moreover, blood PGE_2_ concentration levels were higher in late than early cART responders (74). Thus, PGE immunomodulatory effects during antiviral responses are context-specific and their role in tissue-localized immune responses warrants further investigation.

Herein, we combined multiplex immunofluorescence (mIF), spatial transcriptomics and single cell approaches to comprehensively characterize the F/GC landscape of viremic and cART PLWH. Our data reveal significantly different F/GC cellular and molecular landscapes between Vir and cART LNs and suggest that eicosanoid -related pathways may contribute to both the development of anti-HIV antibody responses and the maintenance of the HIV reservoir.

## Results

### Impaired TFH differentiation profile in cART compared to Vir LNs

We first investigated the F/GC immune landscape across reactive LNs from HIV-negative controls (HIVneg), chronic/viremic (Vir), and anti-retroviral treated (cART) PLWH (**Extended Data Table 1-2)**. LNs from HIVneg individuals exhibited reactive follicular hyperplasia, thus serving as a quality control. We applied a mIF assay that allows for *in-situ* phenotypic analysis of T_FH_ and CD8 T cell subsets based on the expression of 13 markers (**Extended Table 3**, panel1, **Fig.1a and Extended Data Fig.1a**). Identification and quantitative analysis of T_FH_ cell subsets were carried out by Histocytometry (75) focusing mainly on F/GC areas (CD20^dim/hi^ areas, both primary and secondary follicles) (**Fig.1b**). Although lower than in the reactive follicles of the HIV-negative group, comparable cell densities (normalised cell numbers per unit area) of CD3^hi^CD8^lo^PD1^hi^ T_FH_ were found in the Vir and cART groups in this cohort (**Fig.1c**). A significantly higher frequency of less differentiated T_FH_ cells, based on i) the decreased expression of BCL6 and CD57 (10) and ii) the significantly increased prevalence of T_FH_ cells expressing a stem-like TCF1^hi^BCL6^lo^ phenotype, a crucial cell pool for T_FH_ effector responses during chronic viral inflammation (76), was detected in cART compared to Vir F/GCs (**Fig.1d**). In line with this, proliferating T_FH_ (Ki67^hi^) cells were found increased in hyperactive Vir-F/GCs (**Fig.1d**). Furthermore, significantly lower levels of T_FH_ cells expressing surface receptors mediating cell-cell interactions like GITR (77, 78) were found in cART compared to Vir and control LNs while ICOS (79) exhibited comparable levels between the two groups (**Fig.1d and Extended Data Fig.1b**). A TCF1^lo^TIGIT^hi^ cell subset that resembles pre-GC T_FH_ (80) cells was found to be downregulated in cART F/GCs (**Fig.1d**). CXCR3^hi^ T_FH_ cells were enriched in PLWH compared to HIV-negative donors but did not differ between cART and Vir groups (**Extended Data Fig.1b**). Similar profiles were found when the cell densities of the respective cell types were calculated (**Extended Data Fig.1c**). Unsupervised analysis of imaging data using FlowJo plugins (see methods) confirmed the profound heterogeneity of T_FH_ subsets (**Extended Data Fig.1d**). When LN-derived cell suspensions were available, a flow cytometry analysis was applied for some of the markers used for imaging (**Extended Data Fig. 1e**), which further supported the mIF derived data (**Fig.1e**). Therefore, cART favours the accumulation of a less differentiated, stem-like T_FH_ cell subset in PLWH.

**Figure 1.**
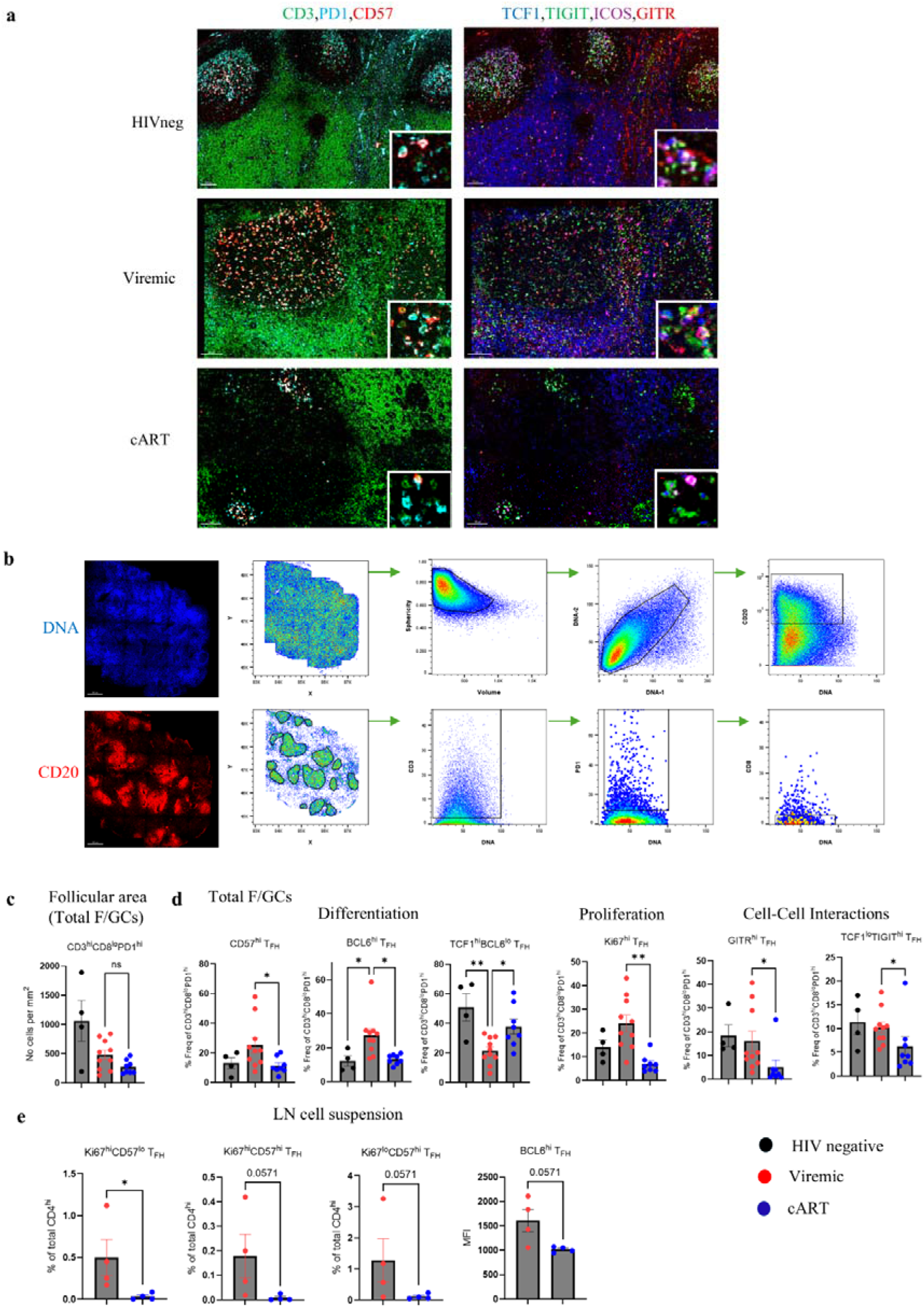
Perturbed TFH differentiation in cART versus Vir lymph nodes. a, Representative mIF images depicting CD57 (red), CD3 (green), PD1(cyan), TIGIT (green), TCF1 (blue), GITR (red) and ICOS (magenta) staining in LNs from control HIV negative individuals, viremic (chronic) and cART PLWH (scale bar: 100 μm). Zoomed areas are shown as insertions at the bottom right of every image. **b,** The Histocytometry gating scheme for the identification of B (CD20^dim/hi^), CD3^hi^ and follicular CD3^hi^PD1^hi^CD8^lo^ cells in a LN is shown. Representative IF images depicting the staining pattern of DNA and CD20 are also included. Individual F/GCs were identified based on the density of CD20^dim/hi^ cells. All follicular areas were combined (boolean gating using FlowJo10 module) for downstream analysis. **c,** Bar graph showing the cell density (normalized per mm^2^) of CD3^hi^CD8^lo^PD1^hi^ T_FH_ cells in LN F/GCs from control HIVneg (black, N=4), Vir (red,N=10) and cART (blue,N=8) donors. Each dot represents a different donor, and bar plots show the mean ± SEM expression. \**P* < 0.05 (Kruskal-Wallis ANOVA test, post-hoc Dunn΄s). **d,** Bar graphs showing the cell frequencies (% of CD3^hi^CD8^lo^PD1^hi^) of CD57^hi^, BCL6^hi^, TCF1^hi^BCL6^lo^, TCF1^lo^TIGIT^hi^, Ki67^hi^ and GITR^hi^, in LN F/GCs from control HIVneg (black, N=4), Vir (red, N=10) and cART (blue, N=8) donors quantified using imaging data via Histocytometry. Each dot represents a different donor, and bar plots show the mean ± SEM expression. \**P* < 0.05 and **P < 0.01.(Kruskal-Wallis ANOVA test, post-hoc Dunn΄s). **e,** Bar graphs showing the cell frequencies (% of CD4^hi^) of Ki67^hi^CD57^lo^, Ki67^hi^CD57^hi^, Ki67^lo^CD57^hi^ and BCL6^hi^ T_FH_ (CXCR5^hi^PD^hi^) in Vir (red, N=4) and cART (blue, N=4) LN cell suspensions samples as measured by Flow Cytometry. Each dot represents a different donor, and bar plots show the mean ± SEM expression. \**P* < 0.05 (Mann-Whitney test).

### Significant reduction of Dark Zone (DZ) GC B cells in cART LNs

Next, the *in-situ* prevalence of B cell subsets was investigated based on relevant phenotypic markers (**Extended Data Table 3**, panel 1 and **Fig.2a**) with Histocytometry analysis (**Fig.2b**). All B cell subsets defined by the expression of BCL6 and Ki67 were found reduced in cART compared to Vir LNs (**Fig.2c**). However, this reduction was statistically significant only for the Ki67^hi^BCL6^hi^CD20^dim^/^hi^ B cells, a subset highly enriched in the DZ (centroblasts) (81) (**Fig.2c**). LN cells *ex vivo* analysis using flow cytometry showed an identical profile (**Fig.2d**). In general, it has been reported that activated T_FH_ and GC-B cells exert reciprocal regulation (79). Although differentiated T_FH_ cell subsets were significantly correlated with the frequencies of Ki67^lo^BCL6^hi^CD20^dim^/^hi^ B cells (enriched in the light zone [LZ]) in Vir LNs, this association was completely lost in cART LNs (**Fig.2e**). Therefore, cART follicular areas are characterized by reduced DZ GC B cells prevalence and uncoupled regulation of T_FH_ and LZ B cells.

**Figure 2.**
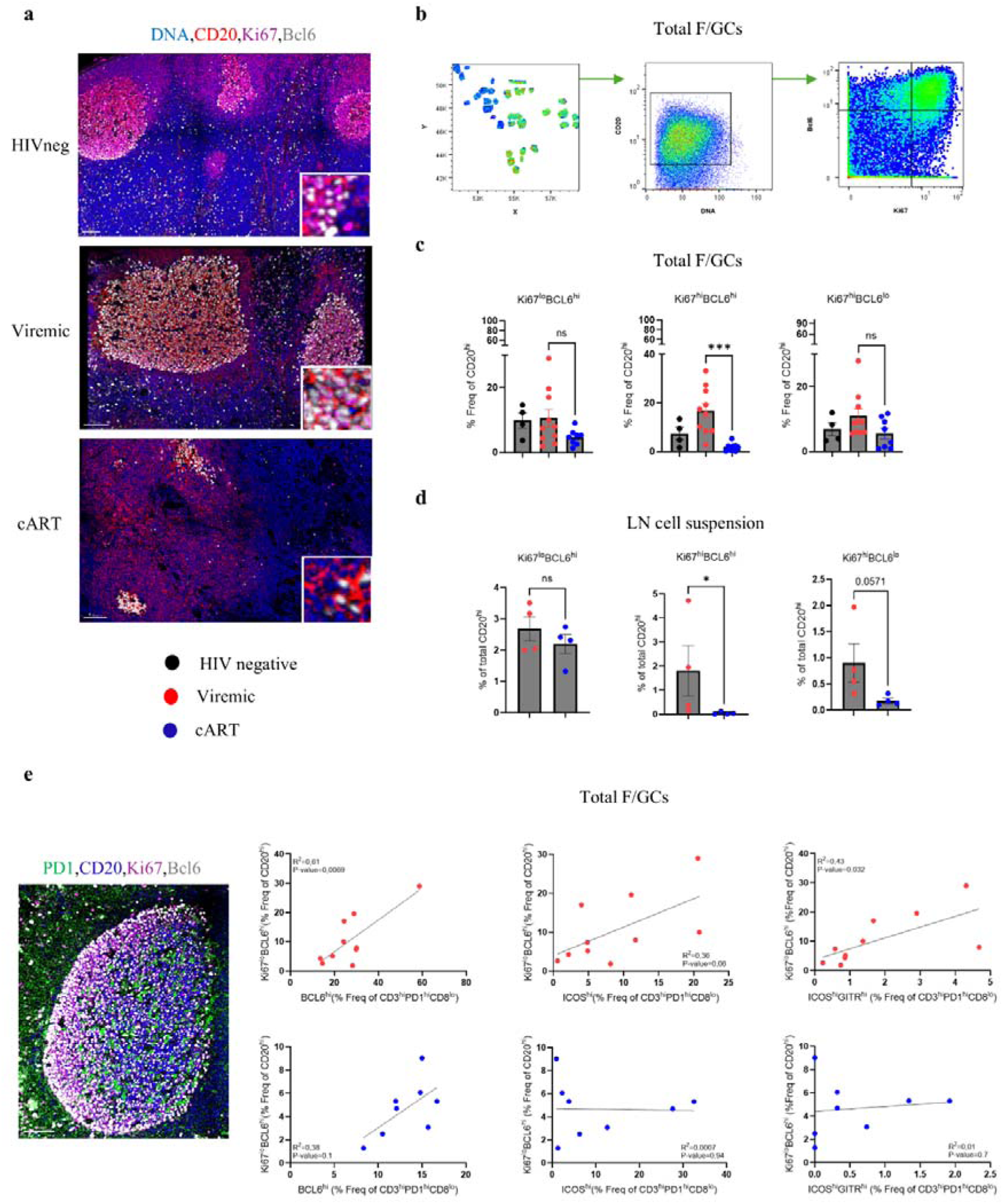
Significant reduction of DZ B cells and loss of reciprocal regulation of LZ B cells with differentiated TFH subsets in cART LNs. a, Representative mIF images depicting CD20 (red), BCL6 (grey), Ki67 (magenta) and DNA/nuclei (blue) staining in LNs from control HIV negative individuals, viremic (chronic) and cART PLWH (scale bar: 100 μm). Zoomed areas are shown as insertions at the bottom right of every image. **b,** The Histocytometry gating scheme for the identification of B (CD20^dim/hi^) and characterization of follicular B cells based on the expression of BCL6 and Ki67 is shown. Individual follicular areas were identified based on the density of CD20^dim/hi^ cells. All follicular areas were combined (boolean gating using FlowJo10) for downstream analysis. **c,** Bar graphs showing the cell frequencies (% of CD20^dim/hi^) of Ki67^hi^BCL6^lo^, Ki67^hi^BCL6^hi^ and Ki67^lo^BCL6^hi^ in LN F/GCs from control HIVneg (black, N=4), Vir (red, N=10) and cART (blue, N=8) donors quantified using imaging data via Histocytometry. Each dot represents a different donor, and bar plots show the mean ± SEM expression. ****P < 0.0001 (Kruskal-Wallis ANOVA test, post-hoc Dunn΄s). **d,** Bar graphs showing the cell frequencies (% of CD20^dim/hi^) of Ki67^hi^BCL6^lo^, Ki67^hi^BCL6^hi^ and Ki67^lo^BCL6^hi^ in Vir (red,N=4) and cART (blue,N=4) LN cell suspensions samples as measured by Flow Cytometry. Each dot represents a different donor, and bar plots show the mean ± SEM expression. \**P* < 0.05 (Mann-Whitney test). **e,** Representative zoomed mIF image of CD20 (blue), BCL6 (grey), Ki67 (magenta) and PD1 (green) depicting a LN follicle from a viremic (chronic) PLWH (scale bar: 100 μm). Linear regression analyses between BCL6^hi^, ICOS^hi^, GITR^hi^ICOS^hi^ T_FH_ and Ki67^lo^BCL6^hi^ B cell frequencies of Vir (red, N=10) and cART (blue, N=8) F/GC cells. R^2^ and p-values are listed. Each dot represents a different donor.

### Significantly reduced effector follicular CD8 T (fCD8) and increased T_FR_ cell prevalence in cART follicles

Given the increased accumulation of ‘effector’ fCD8 T cells in viremic LNs (34) and their possible impact on F/GC cell dynamics and anti-HIV responses (82–84), we sought to characterize the fCD8 T cell profiles in our cohort (**Extended Data Table 3**, panel 1). mIF (**Fig.3a** and **Extended Data Fig.2a**) and Histocytometry analysis (**Extended Data Fig.2b**) showed that despite the comparable frequencies of bulk fCD8 T cells, cART F/GCs harbour significantly lower frequencies of proliferating (Ki67^hi^) and effector (GrzB^hi^) fCD8 T cells compared to Vir LNs (**Fig.3b**), a profile accompanied by significant expansion of the ‘stem-like’ TCF1^hi^ fCD8 T cell compartment (**Fig.3b**). No difference was detected when the trafficking receptor CXCR3 was analysed (**Fig.3b**). Similar profiles were obtained when cell densities were analysed (**Extended Data Fig.2c**). These profiles were not restricted to follicles, as similar patterns were observed in extrafollicular areas (**Extended Data Fig.2d-e**). Flow cytometry analysis showed similar phenotypic profiles in cART compared to Vir LNs either for total CD8 or their follicular (CXCR5^hi^) counterparts (**Fig.3c**).

**Figure 3.**
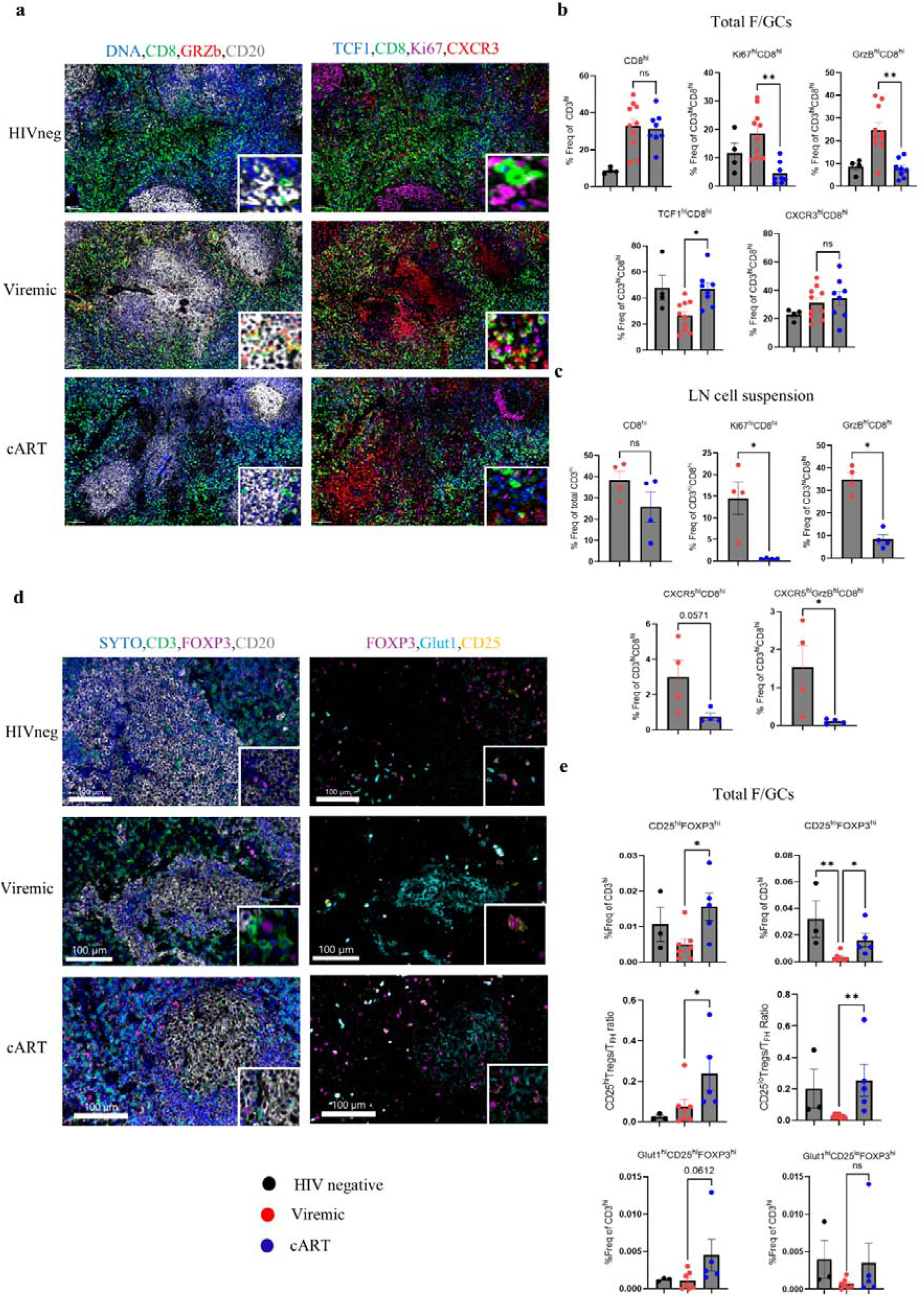
Significant depletion of effector fCD8 cells and enrichment of TFR cells in cART F/GCs a, Representative mIF images depicting CD20 (grey), CD8 (green), GrzB (red), DNA/nuclei (blue), Ki67 (magenta), TCF1 (blue) and CXCR3 (red) staining in LNs from control HIV negative individuals, viremic (chronic) and cART PLWH (scale bar: 100 μm). Zoomed areas are shown as insertions at the upper or bottom right of every image. **b,** Bar graphs showing the cell frequencies (% of CD3^hi^ and CD3^hi^CD8^hi^) of CD8^hi^, Ki67^hi^, GrzB^hi^, TCF1^hi^ and CXCR3^hi^ in in LN F/GCs from control HIVneg (black, N=4), Vir (red, N=10) and cART (blue, N=8) donors quantified using imaging data via Histocytometry. Each dot represents a different donor, and bar plots show the mean ± SEM expression. \**P* < 0.05 and **P < 0.01 (Kruskal-Wallis ANOVA test, post-hoc Dunn΄s). **c,** Bar graphs showing the cell frequencies (% of CD3^hi^ and CD3^hi^CD8^hi^) of CD8^hi^, Ki67^hi^, GrzB^hi^, CXCR5^hi^ and CXCR5^hi^GrzB^hi^ in Vir (red, N=4) and cART (blue, N=4) LN cell suspensions samples as measured by Flow Cytometry. Each dot represents a different donor, and bar plots show the mean ± SEM expression. \**P* < 0.05 (Mann-Whitney test). **d,** Representative mIF images depicting CD20 (grey), CD3 (green), Foxp3 (magenta), DNA/nuclei (blue), Glut1 (cyan) and CD25 (yellow) staining in LNs from control HIV negative individuals, viremic (chronic) and cART PLWH (scale bar: 100 μm). Zoomed areas are shown as insertions at the upper or bottom right of every image. **e,** Bar graphs showing cell frequencies (% of CD3^hi^) of CD25^hi^Foxp3^hi^, CD25^lo^Foxp3^hi^, Glut1^hi^CD25^hi^Foxp3^hi^, Glut1^hi^CD25^lo^Foxp3^hi^ and T_FH_/T_FR_ (both CD25^hi^Foxp3^hi^ and CD25^lo^Foxp3^hi^) cell density ratio in in LN F/GCs from control HIVneg-(black, N=3), Vir (red, N=7) and cART (blue, N=5) donors quantified using imaging data via Histocytometry. Each dot represents a different donor, and bar plots show the mean ± SEM expression. \**P* < 0.05 and **P < 0.01 (Kruskal-Wallis ANOVA test, post-hoc Dunn΄s).

T_FR_ cells, also permissive to HIV infection (85), could represent an additional mechanism of T_FH_ cell regulation in HIV. Thus, we applied a mIF panel (**Extended Data Table 3**, panel 2 and **Fig.3d**) for T_FR_ *in-situ* phenotyping. Glut1, the major glucose transporter for T cells (86), was included as a surrogate marker for their glycolytic capacity. The frequencies of both CD25^hi^ and CD25^lo^ T_FRs_ were increased in cART compared to Vir F/GCs (**Fig.3e**, upper panel). We detected a significantly higher ratio of CD25^hi^FOXP3^hi^ and CD25^lo^FOXP3^hi^ to T_FH_ cells in cART compared to Vir F/GCs (**Fig.3e**, middle panel). Moreover, cART CD25^hi^FOXP3^hi^ cells exhibit substantially increased glycolytic capacity compared to Vir-CD25^hi^FOXP3^hi^ (**Fig.3e**, lower panel). The cell density for both CD25^hi^ and CD25^lo^ T_FRs_ followed the same pattern (**Extended Data Fig.2f**). Flow cytometry data confirmed imaging results showing a trend towards increased CXCR5^hi^FOXP3^hi^ cell frequency and a significantly higher T_FR_/T_FH_ ratio in LNs from cART (**Extended Data Fig.2g**). Overall, our findings suggest that cART is associated with a transition from an effector CD8 T cell-mediated cytotoxic activity to a T_FR_ cell-mediated suppressive microenvironment within follicles.

### Reduced F/GC activity in cART compared to Vir LNs

Follicular innate immune cells like macrophages and dendritic cells regulate F/GC immune responses (87). We analysed the abundance of several innate immune cell types and the *in-situ* expression (bulk cell positivity) of soluble mediators indicating GC activity (CXCL-13, IL-21) by mIF (**Extended Data Table 3**, panel 3) (**Fig.4a and Extended Data Fig.3a-b**). Although the number of visually inspected follicles with detectable FDC-network was comparable between Vir and cART LNs (**Extended Data Fig.3c**), we found higher levels of IL-21 and CXCL-13 either decorated on FDCs or expressed by non-FDC immune cell types in Vir compared to HIVneg and cART LNs (**Fig.4b and Extended Data Fig. 3d**). Interestingly, IL21^hi^CXCL13^hi^ FDC-decorated normalized events were positively correlated with the peripheral blood HIV viral load (logpVL) in viremic PLWH suggesting an association between GC-activity and active HIV transcription (**Extended Data Fig.3e**). Moreover, the reduced activity of cART F/GCs was accompanied by significant reduction of several innate immunity cell types including granulocytes (MPO^hi^) and some macrophage subsets (CD68^hi^CD163^lo^, CD11c^hi^CD68^hi^) in F/GC (**Fig.4c**) and extrafollicular areas (**Extended Data Fig.3f)**. Therefore, impaired F/GC immune responses coincide with reduced innate immune activity in cART F/GCs.

**Figure 4.**
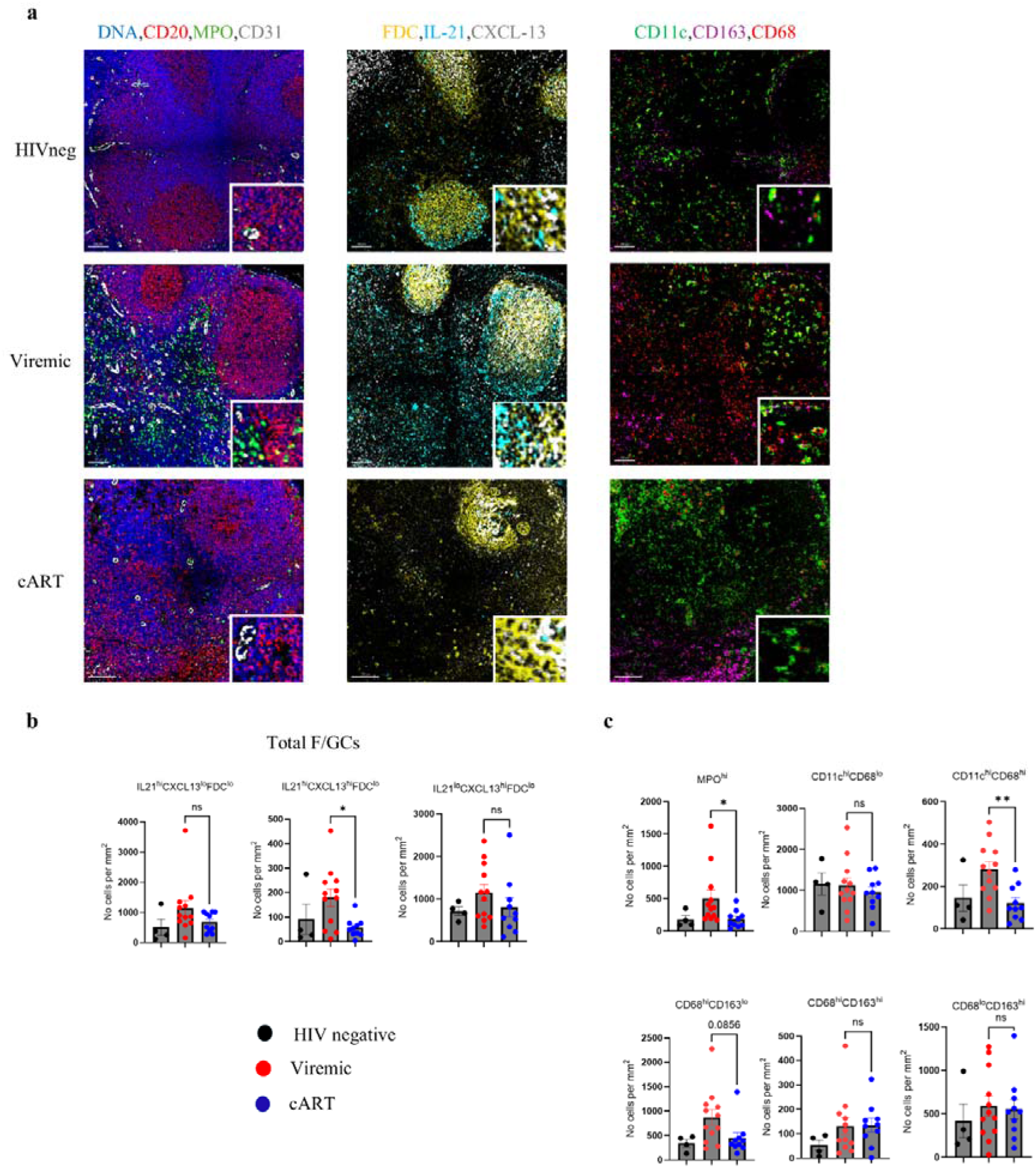
Significant decrease of follicular innate cell subsets and diminished cytokine activity in cART LNs. a, Representative mIF images depicting CD20 (red), MPO (green), CD31 (grey), DNA/nuclei (blue), FDC (yellow), IL-21 (cyan), CXCL-13 (grey), CD163 (magenta), CD11c (green) and CD68 (red) staining in LNs from control HIV negative individuals, viremic (chronic) and cART PLWH (scale bar: 100 μm). Zoomed areas are shown as insertions at the upper or bottom right of every image. **b,** Bar graphs showing the cell densities (normalized per mm^2^) of IL21^hi^CXCL13^lo^FDC^lo^, IL21^hi^CXCL13^hi^FDC^lo^, IL21^lo^CXCL13^hi^FDC^lo^ cells in in LN F/GCs from control HIVneg (black, N=4), Vir (red, N=12) and cART (blue, N=10) donors. Each dot represents a different donor, and bar plots show the mean ± SEM expression. \**P* < 0.05 (Kruskal-Wallis ANOVA test, post-hoc Dunn΄s). **c,** Bar graphs showing the cell densities (normalized per mm^2^) of MPO^hi^, CD68^hi^CD163^lo^, CD68^hi^CD163^hi^, CD68^lo^CD163^hi^, CD11c^hi^ and CD11c^hi^CD68^hi^ cells in in LN F/GCs from control HIVneg (black, N=4), Vir (red, N=12) and cART (blue, N=10) donors. Each dot represents a different donor, and bar plots show the mean ± SEM expression. \**P* < 0.05 **P < 0.01 (Kruskal-Wallis ANOVA test, post-hoc Dunn΄s).

### Downregulation of interferon-and enrichment of AA/Eicosanoid-mediated gene signatures in cART versus Vir follicles

To elucidate the molecular mechanisms underlying the observed alterations, we applied spatial and single-cell (sc) transcriptomic approaches to available tissue material from the same donors (**Extended Data Table 2**). Using the GeoMx platform, we profiled F/GCs as regions of interest (ROIs), defined as CD20-dense areas enriched for CD3^hi^PD1^hi^ cells. Statistical Quantile Learning (SQL) analysis (88) revealed distinct clustering transcriptomic profiles among HIVneg, Vir and cART LN F/GCs (**Fig.5a**). Compared to HIVneg controls, Vir F/GCs were enriched for type I IFN-mediated antiviral and extracellular matrix-related pathways, while cART F/GCs showed enrichment of eicosanoid, IL-10, and IL-12 signalling signatures (**Extended Data Fig.4a-b**). However, key gene pathways associated with the canonical germinal centre reaction, including IL4-, IL-21-and IL1-pathways, were enriched in reactive HIVneg F/GCs (**Extended Data Fig.4a-b**). Differential expression analysis between Vir and cART F/GCs revealed a significant upregulation of interferon stimulated genes (ISGs) and inflammation-related genes in Vir-whereas genes related to antioxidant responses, T-cell homeostasis and endothelial cell function were upregulated in cART F/GCs (**Fig.5b**). Pathway analysis revealed enrichment of pathways related to HIV host interactions, apoptosis/cell cycle pathways, interferon, TCR and BCR signalling in Vir F/GCs confirming thus, the existence of a hyperactive antiviral response (**Extended Data Fig.4c**). Conversely, gene pathways associated with TNF and TGF-b signalling, collagen formation and lipid metabolism were enriched in cART F/GCs (**Extended Data Fig.4c**). Further assessment of the *in situ* operating inflammatory pathways highlighted the upregulation of several interferon signalling-related pathways in Vir-whereas enrichment of arachidonic acid (AA)/Eicosanoid-related gene pathways characterized cART F/GCs (**Fig.5c**). We next curated IFN-and AA/Eicosanoid-gene signatures based on relevant gene pathways and existing literature (55), revealing reduced ISG and increased AA/Eicosanoid-related gene expression in cART-compared with HIVneg and especially Vir F/GCs (**Fig.5d**).

**Figure 5.**
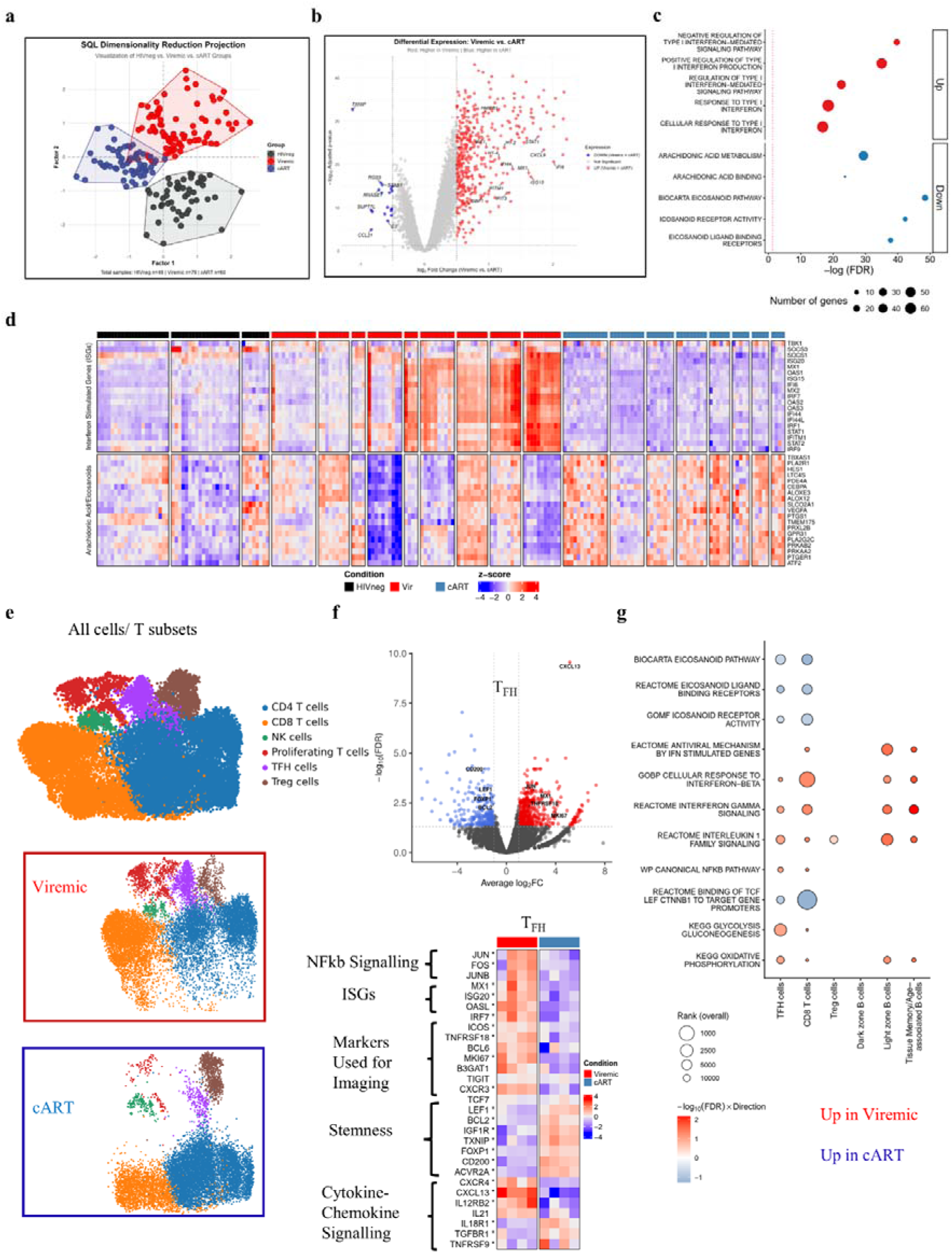
Enrichment of AA/Eicosanoid-related pathways in cART versus Vir follicles. a, SQL clustering projection of HIV-negative (black, N=3, ROIs=49), viremic (red, N=9, ROIs=79) and cART (blue, N=8, ROIs=60) follicular transcriptomic profile. **b,** Volcano plot demonstrating differentially expressed genes between Vir and cART follicular ROIs. Red dots indicate genes upregulated in viremics, blue dots in cART, and grey dots denote non-significant genes. Vertical ticked lines represent a fold-change threshold of 1. **c,** Bubble plot of gene pathways significantly enriched in Vir-(red) versus cART LN F/GCs (blue). Bubble size indicates the number of differentially expressed genes for every pathway. The colour indicates the direction of enrichment (Vir-Red, cART-Blue). The x-axis shows the –log (FDR), corresponding to the negative logarithm of the FDR. **d,** Heatmap displaying differentially expressed genes among all ROIs from HIVneg (black, N=3), Vir (red, N=9) and cART (blue, N=8) (*x*-axis). Genes displayed on the y-axis are classified into functional categories related to IFN/ISGs-and AA/Eicosanoid-signalling. Colour shows z-score transformed counts per million. **e,** UMAP visualization of unsupervised clustering and marker-based cluster annotation of total LN T cells (all cells, Vir and cART, N=8, n=23668), Vir (N=4, n=12880) and cART (N=4, n=10788) as determined by scRNAseq, resulting in 6 different clusters. **f,** Volcano plot of differentially expressed genes between Vir and cART LN T_FH_ (upper panel). Vertical ticked lines indicate the fold-change threshold. Heatmaps displaying differentially expressed genes among T_FH_ (lower panel). Genes displayed on the y-axis are classified into functional categories as noted. Upregulated genes are displayed in red and downregulated genes in blue based on their z-score. **g,** Bubble plot of selected gene pathways significantly enriched in Vir-(red) versus cART follicular ROIS (blue). Gene set enrichment was assessed using limma *fry*. For each cell type, gene sets were ranked by the minimum FDR across the directional and unidirectional tests. Bubble size inversely reflects this rank. Colour indicates the direction of the pathway enrichment and intensity the significance (−log10 FDR), with negative values denoting gene sets up-regulated in cART compared with Vir samples.

Given the limited access to LN cell suspensions, part of Vir (N=4) and cART (N=4) cohorts were analysed using scRNA sequencing. The main bulk cell types of interest B, T, and innate immune cells were annotated by examination of marker gene expression between Leiden clusters (**Extended Data Fig.5a-b**). Clustering analysis revealed considerably fewer cART-derived cells being present in innate cell clusters compared Vir cells, consistent with the mIF data (**Extended Data Fig.5a**). Additionally, alterations in B and T cell subsets were also observed (**Extended Data Fig.5a**). Regarding the B cell compartment, an overall reduction of memory, follicular and plasma cell subsets in cART compared to Vir LNs was detected (**Extended Data Fig.5c-d**). Next, we focused our analysis on the transcriptional profile of T-cell subsets. Relevant T cell subsets were annotated (**Fig.5e** and **Extended Data Fig.6a**) and prominent alterations were revealed between Vir and cART LNs for all identified T cell clusters (**Fig.5e**). With respect to T_FH_ cells, distinct clustering profiles were observed between cART and Vir LNs (**Extended Data Fig.6b**). In line with our imaging analysis, increased gene expression of *TNFRSF18* (GITR), *MKI67* (Ki67), *BCL6, B3GAT1 (*CD57) and *CXCL13/IL21* as well as several ISGs, was found in Vir compared to cART T_FH_ cells (**Fig.5f**). On the other hand, genes indicating a less differentiated profile (*FOXP1*(89)), increased stemness potential (*TCF7/LEF1*(90), *BCL2* (91), *IGF1R* (92), *ACVR2*(93)) and enhanced responsiveness to TGF-beta signalling (*TGFBR1*) were significantly upregulated in cART T_FH_ cells (**Fig.5f**). Regarding CD8 T cells, and in agreement with the imaging results, lower gene expression of *GZMB*, *MKI67* was detected in cART (**Extended Data Fig.6c**). Notably, an enriched activated/exhausted gene profile (*CD38*, *PDCD1*, *TOX2*, *LAG3*) was found in Vir CD8 T cells whereas a less differentiated/stem-cell like (*CCR7*, *IL7R*, *LEF1*, *TCF7*) signature characterized LN CD8 T cells from cART-PLWH (**Extended Data Fig.6c**). Focusing on T_regs_, suppressive capacity-related genes displayed a mixed profile with some being significantly upregulated in Vir (*NR4A2*, *LAG3*, *IDO1*) and others in cART (*ADARB1*, *NFATC3*, *MT-CYB*) T_regs_ suggesting adaptation to distinct tissue microenvironments (**Extended Data Fig.6d**). However, TNF expression, which has been reported to increase T_reg_ proliferation (94), was significantly upregulated in cART T_reg_ (**Extended Data Fig.6d**). Pathway enrichment analysis revealed a strong enrichment of interferon-and IL-1 cytokine signalling in most of the Vir follicular cell subsets and NFkB-related pathways particularly enriched in Vir T_FH_ and CD8 T cells (**Fig.5g**). On the other hand, pathways related to TCF1/LEF1-mediated stemness were particularly enriched in cART T_FH_ and CD8 cell subsets (**Fig.5g**), which further validates our imaging results. Additionally, Vir samples exhibited signs of enhanced metabolism, with increased glycolysis in T_FH_ and CD8 T cells and elevated oxidative phosphorylation in most immune cell subsets (**Fig.5g**). Of note, AA/Eicosanoid signalling pathways were particularly enriched in cART T_FH_ and CD8 cells, implying an enriched prostanoid-mediated immune regulation (**Fig.5g**), consistent with our GeoMx analysis (**Fig.5c**). Therefore, cART is associated with a transition from an interferon-to an AA-Eicosanoid-enriched F/GCs centre, presumably pointing to an altered immunosuppressive microenvironment.

### Distinct spatial distribution of interferon compared to AA/Eicosanoid pathways in F/GCs from PLWH

We extended our spatial biology studies by applying the Visium HD platform, which provides higher resolution (**Extended Data Table 2**). Identification of ROIs (F/GCs) was based on the expression of *MS4A1*/*CD20* (**Extended Data Fig.7a**). Following cell segmentation and ROI identification, downstream analysis was performed on segmented cell data (**Extended Data Fig.7a**). Firstly, we investigated whether the TCF1 and FOXP3 imaging expression profiles align with their *in-situ* mRNA expression (**Fig.6a**). In line with the mIF analysis, a significant difference between Vir and cART F/GCs was found for both the *TCF7* and *FOXP3* expressing T cells (**Fig.6a**). Then, the *in-situ* profile of interferon vs AA/Eicosanoid signatures, using the in-house curated gene sets, was investigated. In line with the GeoMx data (**Fig.5d**), an overall enrichment of the AA/Eicosanoid signature was detected in cART-whereas Vir F/GCs displayed an interferon transcriptomic fingerprint (**Fig.6b and Extended Data Fig.7b**). Given this profile, we next examined whether these molecular signatures were co-enriched *in-situ*. A clear *in situ* segregation was observed between the two signatures when Vir and cART F/GCs were analysed (**Fig.6c**, left panel). Only a small fraction of Vir F/GCs exhibited co-enrichment of both gene signatures (**Fig.6c**). Of note, a similar profile was observed using the GeoMx data (**Fig.6c**, right panel). We next focused on cART F/GCs and found a tendency toward spatial segregation of interferon-and AA/Eicosanoid-enriched areas (**Fig.6d**). Still, a considerable number of the analysed follicles were characterized by dimmer expression of both signatures (**Fig.6d**). To investigate whether an enriched AA/Eicosanoid signature was associated with a distinct transcriptional landscape, we visually stratified cART ROIs according to their respective AA/Eicosanoid gene set scores (**Fig.6d)**. Notably, we observed that AA/Eicosanoid ‘low’ F/GCs were enriched for pathways related to GC-B cell differentiation and isotype switching as well as T cell differentiation (**Fig.6e**, left panel). In contrast, AA/Eicosanoid ‘high’ F/GCs were enriched in pathways associated with eicosanoids and prostaglandins (thus validating our grouping) and pathways associated with cell stemness (**Fig.6e**, left panel). A similar profile was observed when we applied the same analytical pipeline to GeoMx-derived data (**Fig.6e**, right panel). Our data revealed the molecular heterogeneity of follicular regions and further support a role for AA/Eicosanoid signalling in shaping GC-responses during cART.

**Figure 6.**
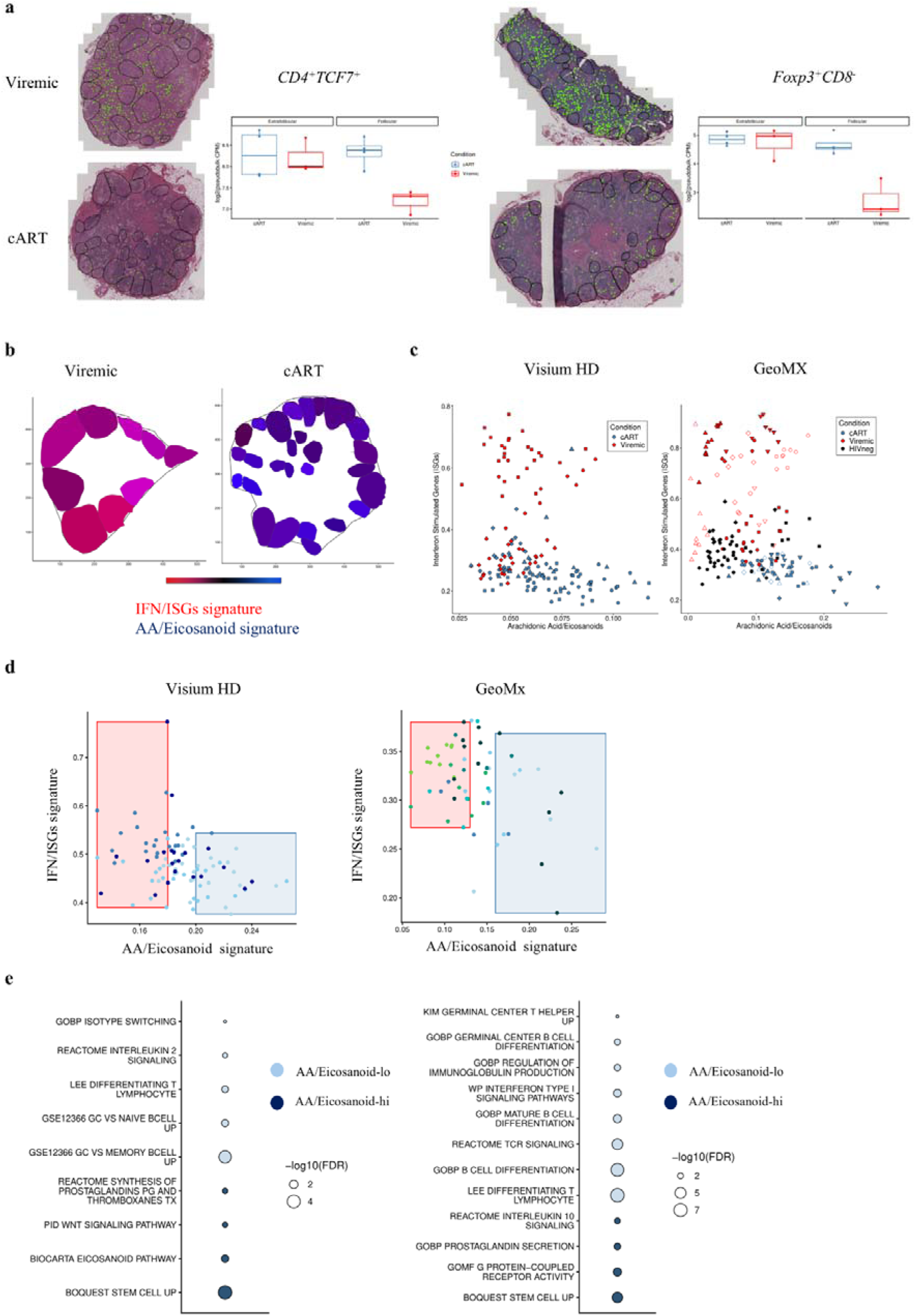
Distinct transcriptional profiles between IFN/ISG-and AA/Eicosanoid-enriched follicular ROIs. a, Representative Space Ranger-derived images depicting cells co-expressing *CD4* and *TCF7* mRNA (left) or expressing *FOXP3* in the absence of *CD8A* and *CD8B* mRNA from viremic (upper panel) and cART LNs (lower panel) of PLWH. Faceted box-and-whisker plots showing pseudobulk log (counts per million (CPM)) in extrafollicular and follicular regions of cART (blue) and viremic LNs (red) are also included. Dots represent individual donors. Black outlines show the convex hull of the follicular annotation based on CD20/MS4A1 mRNA. Green dots are representing segmenting cells expressing the aforementioned mRNAs. **b,** Representative 2D tissue plots demonstrating the spatial gradient of gene signature enrichment across F/GCs from viremic and cART donors, as calculated by UCell. For each signature, the maximum signature score across all samples was mapped to the pure colour (blue or red) and black to zero. The IFN/ISG (red) and AA/eicosanoid (blue) were summed to give the displayed colour values. **c,** Scatter plots demonstrating the Interferon Stimulated Genes (ISG)-and Arachidonic Acid/Eicosanoids-signature scores calculated using either Visium HD (left) or GeoMx (right) data for cART (blue, Visium N=4, GeoMx N=8), Vir (red, Visium N=3, GeoMx N=9) and HIVneg (black, GeoMx N=3) ROIs-F/GCs. Different symbols indicate ROIs from different donors **d,** Scatter plots demonstrating the ISG-and AA/Eicosanoid-signature scores calculated using either Visium HD (left, N=4) or GeoMx (right, N=8) data focusing on cART ROIs. Red and blue boxes are indicative of how ROIs were segregated into AA/Eicosanoid-hi (blue box) and AA-Eicos-low (red box) categories for downstream analyses. Each dot is a follicular ROI colour coded based on donors’ ID. **e,** Dot plot showing significantly enriched gene sets analysed using the limma Camera test. Dot size represents statistical significance (−log FDR), and colour indicates AA/Eicosanoid-hi (dark blue) or AA/Eicosanoid-low (light blue) – enriched pathways.

### In-situ increased expression of PGE receptor EP4 and synthase (mPGES-1) in cART LNs

Based on the above data, we investigated whether eicosanoid pathway-mediators were differentially expressed between Vir and cART follicles. PGE_2_, the most abundant eicosanoid in humans, has been implicated in multiple aspects of HIV pathogenesis (65, 95). PGE_2_ synthesis is tightly regulated by cyclooxygenases (COX) and prostaglandin E synthases (96), while its immunomodulatory effects are mainly mediated through EP2 (moderate sensitivity) and EP4 (increased sensitivity) receptors (65). We sought to investigate and compare the *in-situ* expression of the PGE machinery molecules (**Extended Data Table 3,** panel 4) between Vir and cART LNs using mIF and Histocytometry (**Fig.7a** and **Extended Data Fig.7c**). Albeit not statistically significant, the cell densities of EP4-expressing populations (EP2^hi^EP4^hi^ and EP2^lo^EP4^hi^) displayed a consistent trend towards increased levels in F/GCs and extrafollicular areas of cART compared to Vir LNs (**Fig.7b**). Moreover, although the generic prostaglandin synthesis enzyme COX2 was found to be moderately increased in Vir F/GCs, the PGE -specific enzyme (mPGES-1) was significantly upregulated in both the follicular and extrafollicular areas of cART LNs (**Fig.7c**). Interestingly, a positive correlation was found between the bulk follicular EP2^hi^EP4^hi^ prevalence and the EP4^hi^/ EP2^hi^ ratio with the duration of the pre-ART period (**Fig.7d**). Thus, consistent with the transcriptomic data, *in-situ* imaging revealed increased PGE_2_ sensing and synthesis in cART LNs.

**Figure 7.**
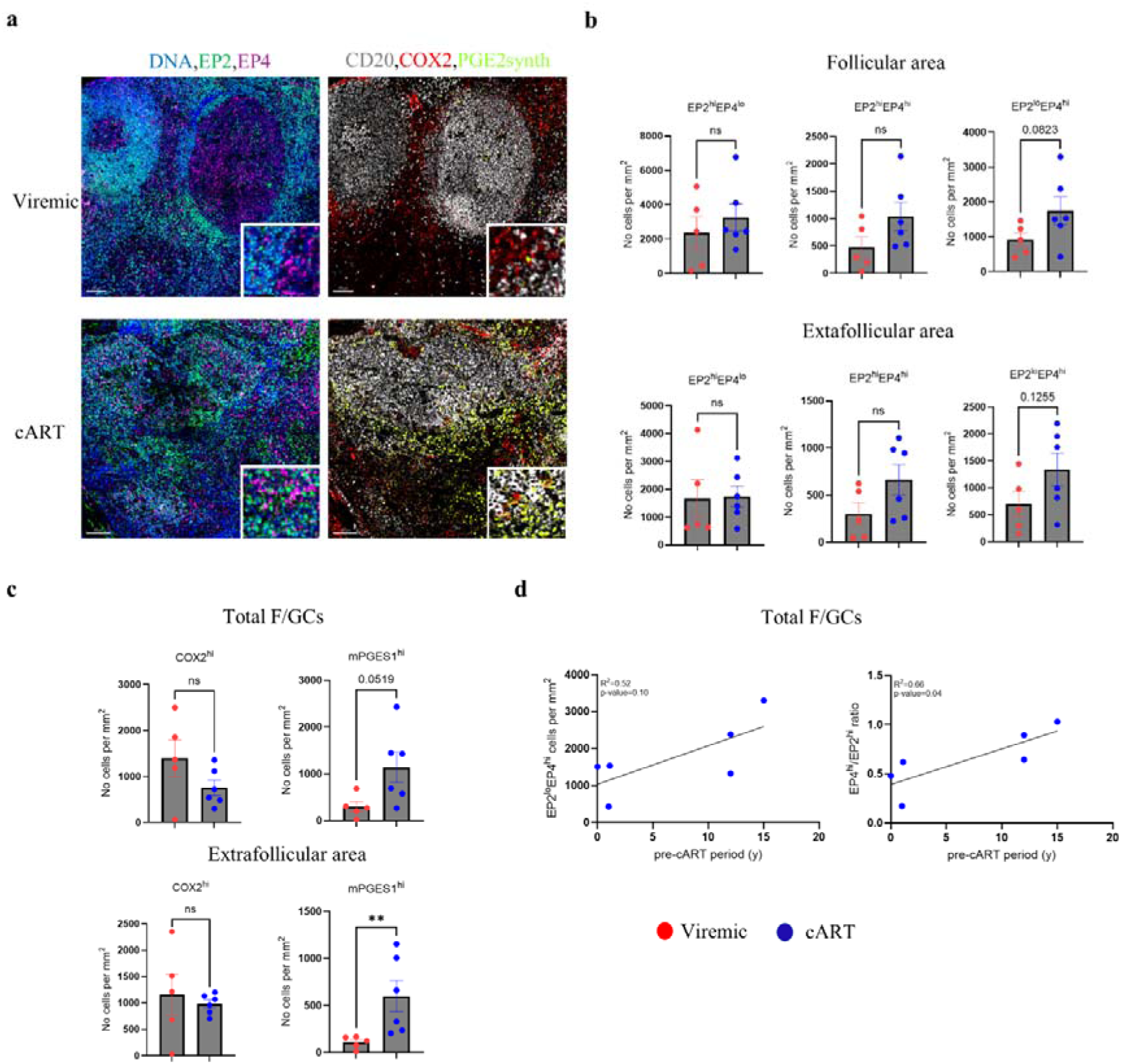
Elevated in situ expression of EP4 and PGE synthase (PGES-1) in cART F/GCs. a, Representative mIF images depicting DNA/nuclei (blue), EP2 (green), EP4 (magenta), CD20 (grey), PGE2 synthase (light green) and COX2 (red) in LNs from viremic (chronic) and cART PLWH (scale bar: 100 μm). Zoomed areas are shown as insertions at the bottom right of every image. **b,** Bar graphs showing the cell densities (normalized per mm^2^) of EP2^hi^EP4^lo^, EP2^hi^EP4^hi^ and EP2^lo^EP4^hi^ from Vir (red, N=5) and cART (blue, N=6) donors in LN F/GCs (upper row) and extrafollicular (lower row) areas. Each dot represents a different donor, and bar plots show the mean ± SEM expression. (Mann-Whitney test). **c,** Bar graphs showing the cell densities (normalized per mm^2^) of COX2^hi^ and mPGES1^hi^ in Vir (red, N=5) and cART (blue, N=6) LN F/GCs (left) and extrafollicular (right) areas. Each dot represents a different donor, and bar plots show the mean ± SEM expression. (Mann-Whitney test) **P < 0.01 (Mann-Whitney test). **d,** Linear regression analyses between EP2^lo^EP4^hi^ cell density or EP4^hi^/EP2^hi^ ratio and the duration (years) of pre-ART period [HIV (years since detection)-cART (years of treatment)] in F/GCs(N=6). R^2^ and p-values are listed. Each dot represents a different donor.

### PGE_2_ as α potential modulator of both antibody responses and viral reservoir in cART LNs

Given the potential suppressive role of PGE_2_ in T_FH_ cell activation and GC immune activity, we asked whether it could also be associated with the development of neutralizing anti-HIV antibodies. To this end, we took advantage of our recent scRNA dataset (15) of LN-derived cells from PLWH with non-neutralizing or broad neutralizing antibodies (**Extended Data Fig.8a)**. Focusing our analysis on type I/II interferon and AA/Eicosanoid pathways we detected an overall higher enrichment of interferon and AA/Eicosanoid signatures in non-neutralizing follicular cell types, particularly in the memory B-cell compartment (**Fig.8a**).

**Figure 8.**
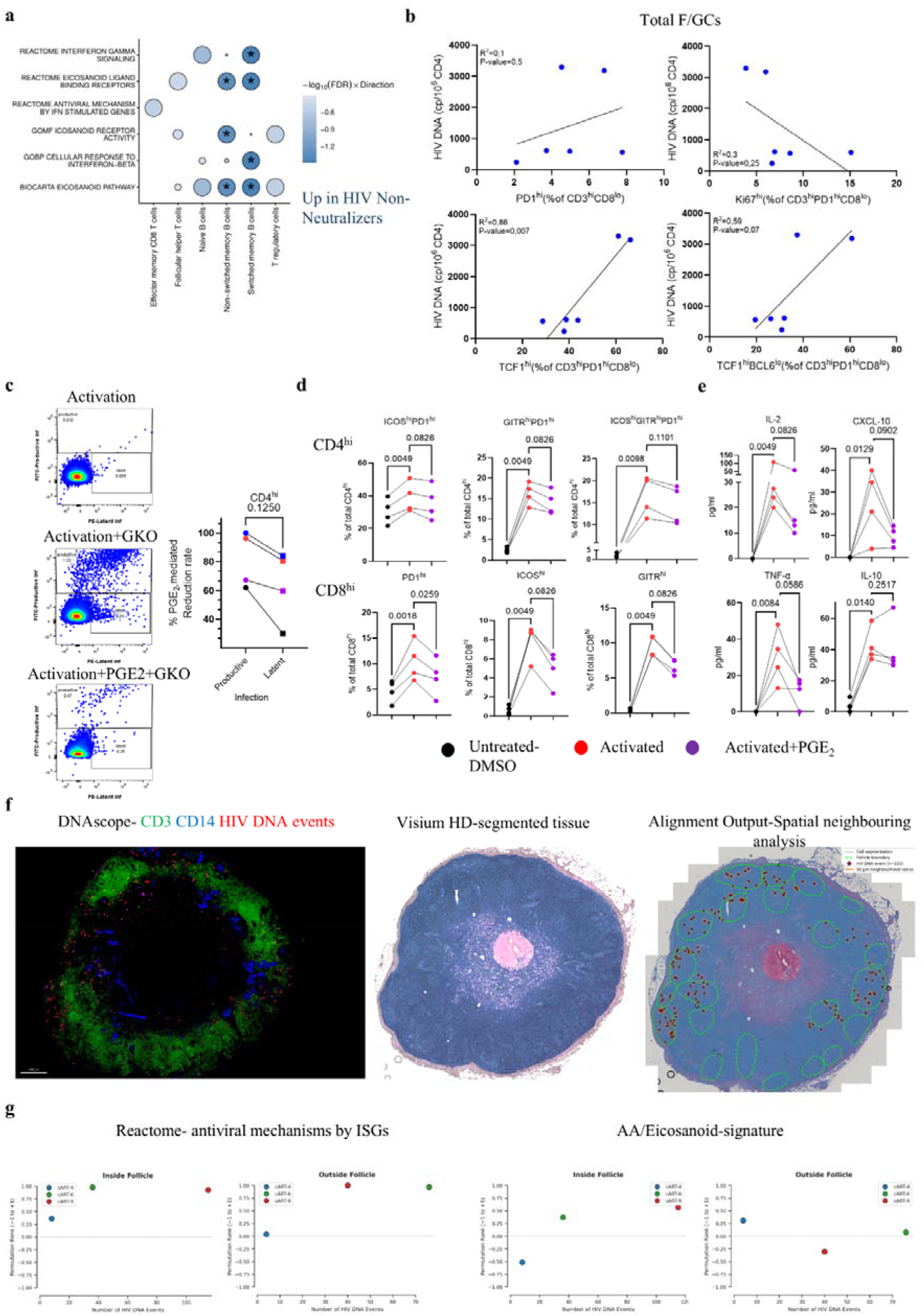
Follicular AA/Eicosanoid enrichment is associated with the absence of anti-HIV neutralizing antibodies in Vir and the maintenance of HIV reservoir in cART PLWH. a, Bubble plot of selected gene pathways significantly enriched in viremic Non-Neutralizers compared with Neutralizers PLWH. Genes set enrichment was assessed using limma fry. Gene sets were ranked by the minimum FDR across tests. Bubble size inversely reflects this rank (larger bubbles = lower rank, more significant). Colour indicates direction and intensity indicates significance (−log10 FDR), with negative values denoting gene sets up-regulated in Non-Neutralizers compared with Neutralizers PLWH. Asterisks are indicative of statistical significance (FDR<0.05). **b,** Linear regression analyses between different T_FH_ subsets (Ki67^hi^, TCF1^hi^ and TCF1^hi^BCL6^lo^) cell frequencies of cART (blue, N=6) LN F/GCs and matched PBMCs HIV DNA measurements (cp/10^6^ CD4 cells). R^2^ and p-values are listed. Each dot represents a different donor. **c,** Flow cytometry histoplots demonstrating the identification of *in vitro* Productively (FITC) - and Latently (PE)-GKO infected cells upon activation with or without PGE2 administration. Graph with connecting lines demonstrating the % PGE_2_-mediated Reduction rate between Productively (circles) and Latently (squares) infected cells. Different donors are colour coded (N=4). **d,** Graphs showing human primary tonsillar cell frequency (%) of control (DMSO) and activated, with or without concomitant PGE_2_ administration, CD4 and CD8 subsets based on PD1, ICOS and GITR expression (N=4). P-values are also shown. *P < 0.05 **P < 0.01 ***P < 0.001 (one-way ANOVA tests). **e,** Graphs showing the concentration of IL-2, CXCL-10, TNF-α and IL-10 in supernatants from control (DMSO) and activated, with or without concomitant PGE_2_ administration, human primary tonsillar cells (N=4). P-values are also shown. *P < 0.05 **P < 0.01 (Friedman one-way ANOVA tests). **f,** Representative cartoon to describe the multimodal integration of HIV DNA scope (mIF image showing CD14 (blue), CD3 (green), HIV-DNA events (red)) to Visium HD (segmented cells in blue). HIV DNA events were represented, based on their x,y coordinates) as Imaris-generated spheres to help with their visualization. The different methods were applied in sequential or proximal sections. The result demonstrates the result of our *in-silico* alignment. **g,** Number of HIV DNA events detected inside F/GCs (left) and in the extrafollicular area (right) versus permutation rank for cART-4 (blue), cART-6 (green), and cART-9 (red). Positive permutation rank values indicate enrichment of the relevant gene pathways (Antiviral mechanisms by ISGs and AA/Eicosanoid gene sets) relative to the null distribution, while values near zero (denoted with a dashed horizontal line) indicate no enrichment.

T_FH_ are well-established HIV-reservoirs (31). For part of the cART samples, matched PBMCs were also available. Analysis of cell-associated HIV-DNA revealed a positive correlation with the *in-situ* F/GC abundance of TCF1^hi^ T_FH_ cells while a trend for negative association with Ki67^hi^ T_FH_ cells was found (**Fig.8b**). Of note, a positive association was also observed between peripheral HIV-DNA latently infected and follicular EP2^hi^EP4^hi^ cells (**Extended Data Fig.8b**). To investigate a potential role of PGE_2_ in HIV latency, *in vitro* infection experiments were carried out using a dual reporter virus (97) and activated CD4^hi^ T cells from HIVneg donors in the presence or not of PGE_2_ (**Fig.8c** and **Extended Data Fig.8c**). Although viral infection was overall reduced in the presence of PGE_2_, this reduction was consistently higher in ‘productively’ compared to ‘latently’ infected, activated CD4 T cells (**Fig.8c**). Further assessment of its potential role showed that PGE_2_ reduces *in vitro* activation of both CD4 and CD8 T cells as shown by the decreased expression of ICOS, GITR and PD1 (**Fig.8d** and **Extended Data Fig.8d**). Moreover, the secretion of several Th_1_ cytokines and chemokines including TNFα, CXCL-10 and IL-2 was also reduced (**Fig.8e** and **Extended Data Fig.8e**).

We extended our spatial biology studies by performing an *in-silico* alignment of Visium HD and HIV DNAscope generated images from proximal tissue sections (**Fig.8f** and **Extended Data Fig.9)**. Despite the small number of tissues analysed (n=3), a positive correlation between the PBMCs HIV DNA and the *in situ* calculated number of LN HIV DNA^+^ cells was detected (**Extended Data Fig.10a).** Integration of DNAscope and Visium HD data provided insights into the transcriptional microenvironment surrounding HIV-DNA^+^ CD3^hi^ T cells. Thus, a spatial neighbouring analysis to assess the potential *in-situ* enrichment of interferon and AA/Eicosanoid gene signatures around (radii=50 μm) HIV DNA^+^ CD3^hi^ T cells was conducted by applying a permutation analytical pipeline that also assesses the randomness of the observed profiles (see Methods and **Extended Data Fig.10b**). With respect to publicly retrieved interferon-related gene signatures (both type I and II), a similar profile between follicular and extrafollicular areas was found with increasing probability for non-random enrichment of the signatures which were positively associated with the number of HIV DNA^+^ CD3^hi^ cells (**Fig.8g, Extended Data Fig.10c-d**). The compact in-house IFN signature that we curated did not recapitulate the same pattern, potentially reflecting its limited gene set (**Extended Data Fig.10d).** Regarding the AA/Eicosanoid signature, a similar profile, with that of publicly available IFN-gene sets, was observed only for the follicular areas, although at lower magnitude of significance for non-random enrichment (**Fig.8g and Extended Data Fig. 10c**). Our data point to AA/Eicosanoids, and particularly PGE_2_, as a potent regulator of F/GC immune responses with possible impact on neutralizing antibody generation and the maintenance of follicular viral reservoir.

## Discussion

Comparison of Vir and cART F/GCs revealed tissue signatures associated with inflammatory states and active HIV transcription. The consistency of our findings across complementary *in-situ* and single-cell approaches validated that the observed profiles reflect robust tissue-level features rather than section-specific variability.

Imaging analysis revealed a less differentiated phenotype of T_FH_ cells in cART compared to Vir F/GCs. In line, scRNA analysis revealed significantly higher expression of genes favouring stemness and lower expression of genes favouring differentiation of T_FH_ cells in cART compared to Vir PLWH. This T_FH_ profile was associated with a reduction of the DZ-B compartment, evidence of dysregulated GC activity, and with the concomitant disconnection of the proposed T_FH_ / GC B cells mutual regulation (79) in cART LNs. The cART T_FH_ cell profile may reflect intrinsic characteristics or an altered microenvironment shaped by immunoregulatory cells, suppressive tissue factors, and low-grade inflammation. fCD8 T cells and T_FR_ cells represent two major regulators of F/GC immune dynamics in HIV infection (34, 45, 47, 98). Similarly to T_FH_ cells, fCD8 T cells exhibited a less differentiated profile in cART compared to Vir LNs, suggesting a potential role for shared microenvironmental factors. T_FR_-mediated activity can regulate the effector functions of both fCD8 and T_FH_ (99, 100). The higher frequency of potentially glycolytic T_FRs_ in cART suggests that in addition to their abundant presence, their suppressive function could be also enhanced extrinsically in cART compared to Vir LNs. The metabolic switch of T_regs_ and T_FRs_ from oxidative phosphorylation to glycolysis has been associated with non-canonical suppressive activity (101, 102). Therefore, T_FRs_ - mediated immunosuppressive activity, together with reduced CD8 cytotoxicity, create an unfavourable microenvironment for anti-HIV responses in cART F/GCs.

As expected (103, 104), cART was associated with a downregulation of LN inflammation, judged by the overall reduced prevalence of several proinflammatory innate immune cell populations, including MPO^hi^ neutrophils and macrophage subsets (105, 106), in cART LNs. The observed low-grade inflammation is associated with significant reduction in IL-21 and CXCL-13 production, two soluble mediators with critical impact on GC development. Our spatial transcriptomic analysis showed a significant overexpression of signals mediating GC development (e.g. IL-21, IL-4, IL-1 signalling) in follicles from HIVneg compared with cART LNs, further pointing to multifactorial impaired GC response in the latter. Our data suggest that the low-inflammatory, T_FR_-enriched cART F/GC microenvironment may impair T_FH_ differentiation and T_FH_–GC B cell interactions, potentially compromising B cell responses in PLWH.

Spatial biology analysis clearly showed a shift from a potentially tissue-damaging, interferon-mediated inflammation in Vir F/GCs to a presumably less immunopathological and tolerogenic AA/Eicosanoid-mediated inflammatory microenvironment in cART F/GCs. scRNA analysis extended these observations by revealing a significant enrichment of interferon fingerprints across GC B and T cell subsets in Vir compared to cART LNs. By contrast, cART-treated LNs were characterized by a significantly enriched AA/Eicosanoid signature in T_FH_ and CD8 T cells specifically, suggesting a more restricted, cell-specific action of this pathway as a regulator of F/GC activity. Therefore, in addition to T_FR_ cells, the suppressive activity within the cART F/GC microenvironment could be further augmented by the local operation of AA/Eicosanoids, particularly PGE_2_, a central mediator of this pathway (65). PGE_2_ has a negative impact on CD8 activation (67), regulates T_FH_ differentiation (107) and can strengthen T_reg_ suppressive capacity (108). In contrast to leukotrienes such as LTA4, prostaglandins like PGE_2_ suppress i) interferon production (69, 109) and ii) Th1 cytokines’ signalling (e.g IL-2) while promoting Th2, Th17 and T_reg_ responses (65, 110–112). In line with these reports, we found that *in-vitro* PGE_2_ treatment i) suppressed tonsillar CD4/T_FH_ and CD8 T cell activation and ii) reduced the secretion of Th1 cytokines/chemokine (IL-2, TNF-α, IP-10), whereas suppressive (IL-10) or other proinflammatory (IL-6) cytokines were minimally affected. Therefore, eicosanoids such as PGE_2_ could act as molecular coordinators of direct and indirect (i.e. through fostering T_FR_ cell activity) suppressive effects on T_FH_ and CD8 T cells in cART LNs.

Interferons tend to suppress the processing and signalling of AA (65, 113, 114). However, a positive regulation has also been demonstrated (115), suggesting that the outcome of the interaction between these two pathways is cell type-and context-dependent. The contrasting expression patterns of interferon-and AA/Eicosanoid-related genes (GeoMx analysis), together with the limited co-enrichment of their respective signatures within Vir and cART F/GCs (Visium HD and GeoMx analyses), indicate compartmentalization or reciprocal regulation between these biological programs in PLWH LNs.

F/GC adaptive immune cells are of great importance for the development of effective pathogen-and immunogen-antibody responses as well as the maintenance of the HIV reservoir. scRNA analysis of an independent viremic PLWH cohort revealed enriched interferon and AA/Eicosanoid signatures, particularly in B cells, in LNs lacking neutralizing activity compared with those with neutralizing activity. We hypothesize that these lipid mediators could act as potent negative regulators of humoral responses in cART PLWH, acting directly on T_FH_ and B cells or indirectly (e.g. through T_FR_ cells). *In-vivo* blockade of PGE_2_ signalling (e.g. using EP2/4 blockers (116, 117) in preclinical models could directly validate its suppressive impact on GC responses and assess its potential to enhance T_FH_ activity by removing a regulatory brake. Given the promising outcomes of COX2-mediated prostaglandin synthesis inhibition in PLWH receiving ART (72, 118) targeted inhibition PGES-1 may represent a more selective strategy to further enhance anti-HIV responses. This approach will lead to PGE2-specific blockade without affecting the production of other potentially beneficial COX2-related prostaglandins that exhibit immune-boosting properties.

Despite the limited sample size, our data suggest that the accumulation of stem-like T_FH_ cells in cART LNs may support HIV latency through their stable kinetics, even though the T_FH_ cell pool is reduced after long-term therapy (119). Long-lived TCF1^hi^ T_FH_ cells are characterized by elevated expression of cAMP-regulated genes which could indicate sustained PGE_2_-signalling (120). *In vitro* infection analysis showed a weaker impact of PGE_2_ on latent HIV DNA accumulation than on the actively transcribed viral pool. The integrative multimodal analysis combining Visium HD and mIF HIV DNAscope provides a novel *in-situ* approach for the comprehensive characterization of the viral reservoir microenvironment. Our data suggest that in addition to interferon-mediated antiviral function (121), AA/Eicosanoids may contribute to the viral reservoir maintenance within F/GCs. Given i) the cART-mediated suppression of *in-situ* interferon signature and the potential sustained AA/Eicosanoid signalling in long-term treated PLWH and ii) the spatial segregation of interferon and AA/Eicosanoid signals, with enhanced AA/Eicosanoid-mediated signalling in the absence of interferon signals, we propose that AA/Eicosanoids may shape the HIV reservoir microenvironment under conditions of limited interferon-mediated antiviral activity. However, this hypothesis requires validation in larger, independent cohorts.

Overall, our data suggest that AA/Eicosanoids, particularly PGE_2_, could represent molecular mediators that coordinate a suppressive, low-grade inflammatory microenvironment. This has a negative impact on the generation of effective B-cell responses and a potential effect on maintaining the HIV reservoir by favouring the presence of stem-like T_FH_ in cART PLWH. Additionally, enhancing anti-HIV CD8 T cell responses during ATI, combined with inhibition of suppressive signals such as eicosanoids, may improve control of viral rebound (122).

Our cross-sectional study cannot establish the chronological sequence of the observed cellular and molecular dynamics. Longitudinal LN sampling in non-human primate SIV models could further validate these findings and mechanistically investigate them. Although we focused on PGE_2_, as an immunomodulatory mediator of LN anti-HIV responses, we cannot rule out effects from other AA-derived molecules. Additionally, in the absence of LN cell measurements, we examined the correlation between HIV latency and the accumulation of LN cell subsets using blood HIV DNA measurements. Previous studies have reported a positive correlation for these two tissue measurements (blood vs LN) at least for PLWH with high latent HIV DNA levels (123). Despite these limitations, our study provides, to our knowledge, the first comprehensive cellular and molecular characterization of the F/GC landscape across untreated and treated-PLWH.

## Methods

### Human Material

The tissue samples included in this study (**Extended Data Table 1**) were obtained from (i) the Centro de Investigacion en Enfermedades Infecciosas (CIENI) (Ethics Committee, CONBIOETICA-09CEI-003-20160427, protocol B03-16), Instituto Nacional de Enfermedades Respiratorias, Mexico City, Mexico (viremic LNs), (ii) the University of Washington (study ID: STUDY00001091, CR ID:CR00004042), Seattle, WA, USA (cART HIV LNs) and (iii) the biobank of the Institute of Pathology of Lausanne University Hospital, Switzerland (HIV negative control LNs). Their use was approved by the Ethical Committee of the Canton de Vaud, Switzerland (protocol number 2021-01161). Tonsillar cell suspensions were obtained from anonymized children who underwent routine tonsillectomy at the Hospital de l’Enfance of Lausanne (protocol, PB_2016-02436 (201/11)). Human Peripheral Blood Mononuclear cells (PBMCs) were obtained from anonymized adult donors (Service of Immunology and Allergies, CHUV). More specifically, blood samples were obtained at the local blood bank (Centre de transfusion sanguine (CTS), Lausanne, Switzerland). The use of samples used in this study was approved by the Institutional Review Board of the CTS, and all subjects gave written informed consent. Only individuals with no sign of HIV, HAV, HBV and HCV infections were included. All procedures complied with the Declaration of Helsinki and received approval from the appropriate Institutional Review Board/Ethical Committee. All tissue samples from PLWH were procured with explicit written informed consent from participants prior to donation, in full accordance with the principles outlined in the Declaration of Helsinki. Samples were used for each method based on tissue and LN suspension availability (**Extended Data Table 2**). Details about the gender of the participants in the study are reported in **Extended Data l Table 1**. Gender was not investigated as a biological variable in our study and mirrors the gender distribution in the clinical cohort from which the participants were drawn. Future work is needed to also extend our findings in females.

### Tissue Processing and staining

Routinely processed formalin-fixed, paraffin-embedded (FFPE) blocks were cut into 5 μm sections and placed on Superfrost glass slides (Thermo Scientific, Waltham, MA, USA, Ref. J1800AMNZ), dried overnight and stored at 4 °C. Before staining, the slides were heated on a metal hotplate (Stretching Table, Medite, Burgdorf, OTS 40.2025, Ref. 9064740715) at 65 °C for 20-30 min. Tissue sections were stained with titrated antibodies using a Ventana Discovery Ultra Autostainer (Roche Diagnostics, Ventana Medical Systems, Tucson, AZ, 85755, USA). Tissues were deparaffinized, rehydrated and the protein epitopes were retrieved using the standard Ventana Discovery’s protocols. Before each antibody incubation step, tissues were blocked using Antibody Diluent/Block from Akoya (ARD1001EA, Akoya Biosciences, Marlborough, MA 01752, USA). The cycling staining/imaging approach that we developed is divided in two different cycles (**Extended Data Table 3**) and the pipeline was thoroughly described in (14). The antibodies used for the different imaging panels are listed in **Extended Data Table 3**.

### GeoMX spatial transcriptomics

Transcriptomic profiling was performed using the commercially available platform GeoMx Digital Spatial Profiling (NanoString) according to the manufacturer’s instructions. FFPE LN sections from viremic PLWH (N=8), cART PLWH (N=7) and HIVneg controls (N=3) were used. Follicular Regions of Interest (ROIs) were identified based on the CD3, CD20, PD-1 *in-situ* staining pattern (active follicles, characterized as CD20-dense areas populated by CD3^hi^PD1^hi^ cells were selected for analysis) before the probe-hybridization step. For data analysis, quality control (QC) and batch correction previously described workflows were applied, following the recommendations in the R package standR (124). We used the Statistical Quantile Learning (SQL) method, a tool for nonlinear dimensionality reduction, to visualize spatial transcriptomic data from HIVneg, Vir and cART F/GCs. Unlike local methods (e.g., UMAP), SQL learns a global latent space via a smooth generator function, enabling interpretable, statistically grounded group separation (https://github.com/jbodelet/SQL). Differential expression between HIVneg, viremic, and cART groups was assessed using limma-Voom (125) with offsets calculated by RUV included to account for patient-specific effects. Enrichment analysis (GSEA) on Reactome pathways was performed using limma Fry method. Gene set tests were performed using limma Fry (126), using all gene set collections except the MIR set from the msigdb package.

### VISIUM HD spatial transcriptomics

For each FFPE block, one freshly cut section of 5 μm-thick was placed onto a Superfrost+ adhesion slide (Epredia). Slides were then heated at 42 °C for 3 h on a thermocycler using the Low-Profile Thermocycler Adaptor (10XGenomics). FFPE slides that were not processed directly were stored at 4°C up to 2 weeks. Deparaffinization, H&E staining, Imaging and Decrosslinking were performed following the 10X Genomics protocol “Visium HD FFPE tissue preparation handbook” (CG000684). Images were acquired with the slide scanner Axioscan 7 (Zeiss). Tissue transfer was done with the CytAssist machine and probe-based libraries were constructed according to the protocol “Visium HD Spatial Gene Expression Reagent kits user guide” (CG000685). The quality control of resulting libraries was performed with the Fragment Analyzer (Agilent Technologies) and the Qubit High-Sensitivity dsDNA assay kit (Invitrogen). Sequencing was performed on AVITI (Element Biosciences) following 10X parameters recommendation (Paired-end, dual indexing: Read 1: 43 cycles; i7 Index: 10 cycles; i5 Index: 10cycles; Read 2S: 50 cycles).

Loupe Browser v9.0 was used to manually align H&E images with Visium CytAssist images. Then, Space Ranger v4.0 was used to map FASTQ files to the 10x Genomics 2024-A human reference, perform cell segmentation and aggregate reads to cells using the Loupe alignments. When multiple tissue pieces were sequenced on a single Visium slide capture region, the alignment and quantification pipeline was performed separately for each tissue piece. Follicles were manually annotated in Loupe Browser by visualising CD20/*MS4A1* gene expression overlayed on the microscopy images. Low per-cell read coverage made cell-level quantitative analyses unfeasible, particularly for the cART samples. All differential expression and gene set enrichment analyses were therefore performed on read counts pseudobulked per follicle. Cell-level expression is shown only for visualisation. Differential expression was performed using R (v4.5.2) edgeR (v4.8.2) with donor included as a blocking factor. Gene set enrichment analyses were performed using limma Fry (v3.66.0) with all gene sets from the msigdb collections except for the MIR set. UCell (v2.14.0) was used to calculate scores for gene set signatures across counts pseudobulked across follicles, using the top 5000 most highly expressed genes.

### sc-Multiome Gene expression

Following thawing, 1 × 10 cells were transferred into PBS containing 0.04% bovine serum albumin (BSA) and processed for DNase treatment and nuclei isolation according to the 10x Genomics Nuclei Isolation for Single Cell Multiome ATAC + Gene Expression protocol (CG000365, Rev. C; see below). Lymph node mononuclear cell (LNMC) nuclei were isolated according to the manufacturer’s instructions (10x Genomics Nuclei Isolation for Single Cell Multiome ATAC + Gene Expression protocol; CG000365, Rev. C). Single-cell gene expression (GEX) libraries were generated using the Chromium Single Cell Multiome Gene Expression platform (10x Genomics). For each sample, 10,000 nuclei were loaded. Isolated nuclei were transposed and partitioned into Gel Beads-in-emulsion (GEMs) using the Chromium Controller and Next GEM Chip J.GEX libraries were generated from the same pool of pre-amplified transposed DNA and cDNA and following the Chromium Next GEM Single Cell Multiome ATAC + Gene Expression User Guide (CG000338). Library quality and fragment size distributions were assessed using the Agilent Bioanalyzer High Sensitivity DNA assay (Agilent Technologies). GEX libraries were sequenced on Illumina NovaSeq S4 flow cells, using 10x Genomics–recommended sequencing parameters. The ATACseq data were not used for the scope of this study.

### Statistics

*In-situ*, *ex-vivo*, and *in-vitro* data were analysed using non-parametric paired (Wilcoxon signed-rank) or unpaired (Mann–Whitney U) tests. For comparisons involving more than two groups, one-way parametric or non-parametric ANOVA (Friedman or Kruskal–Wallis tests, respectively) was performed, followed by the appropriate post hoc tests. Regression analyses were performed to assess associations between variables. Graphs were generated using GraphPad Prism 10.4.1, and statistical significance was defined as p < 0.05.

## Additional methodological details are provided in the Extended Data file

### Lead contact

Further information and any requests should be directed to and will be fulfilled by the lead contact, Konstantinos Petrovas.

## Materials availability

This study did not generate new unique reagents.

## Data and code availability

Data reported in this paper and any additional information required for reanalysis are available from the lead contact upon request. The authors agree to share all publication-related data. For further information, please contact the corresponding author at.

## Acknowledgements

The authors would like to thank Dr Natalie Piazzon, Damien Maison (Tissue Biobank), and Emilie Lingre, Institute of Pathology, CHUV, for their help with tissue processing, Do Rosario Nadine for performing the Luminex Multiplex Cytokine assay as well as Fahr Noemie and Le Gal Tania (Dangaj Lab,UNIL), who assisted with the VISIUM HD experiments.

## Authors Contributions

S.G. design and performance of experiments, data asquisition and analysis, data interpretation, drafting of manuscript. H.L. scRNA and spatial transcriptomics.data analysis and interpretation, drafting of manuscript. M.O. design and performance of mIF experiments, data asquisition and analysis, J.L.R. scRNA and spatial transcriptomics.data analysis and interpretation. P.M.D.R.E. Flow cytometry data acquisition and analysis performed scRNA experiments. B.R. *in vitro* infection experiments, data acquisition and analysis. C.B and S.K assisted with spatial transcriptomic analysis. F.T.R., Y.A.L.V., G.S.M.O., and L.D.L. provided participant material and clinical data. M.P. supervised *in vitro* infection experiments and analysis. M.D performed HIV DNA measurements and analysis. O.L drafted the manuscript. D.D.L supervised VISIU HD analysis, drafted the manuscript. L.V. supervised HIV DNA measurements and analysis, drafted the manuscript. S.P.R. supervised flow cytometry experiments and collection of scRNA data, drafted the manuscript. R.G. supervised scRNA and spatial transcriptomics data analysis, drafted the manuscript. C.P. conceived, designed and supervised the study, interpreted data, drafted and revised the manuscript.

## References

1. Cohen MS, Chen YQ, McCauley M, Gamble T, Hosseinipour MC, Kumarasamy N, et al. Antiretroviral Therapy for the Prevention of HIV-1 Transmission. N Engl J Med. 2016;375(9):830–9.

2. Palmer S, Josefsson L, Coffin JM. HIV reservoirs and the possibility of a cure for HIV infection. J Intern Med. 2011;270(6):550–60.

3. Coiras M, López-Huertas MR, Pérez-Olmeda M, Alcamí J. Understanding HIV-1 latency provides clues for the eradication of long-term reservoirs. Nat Rev Microbiol. 2009;7(11):798–812.

4. Oxenius A, Price DA, Günthard HF, Dawson SJ, Fagard C, Perrin L, et al. Stimulation of HIV-specific cellular immunity by structured treatment interruption fails to enhance viral control in chronic HIV infection. Proc Natl Acad Sci U S A. 2002;99(21):13747–52.

5. Tortellini E, Fosso Ngangue YC, Dominelli F, Guardiani M, Falvino C, Mengoni F, et al. Immunogenicity and Efficacy of Vaccination in People Living with Human Immunodeficiency Virus. Viruses. 2023;15(9).

6. Qi H, Kastenmüller W, Germain RN. Spatiotemporal basis of innate and adaptive immunity in secondary lymphoid tissue. Annu Rev Cell Dev Biol. 2014;30:141–67.

7. Dimopoulos Y, Moysi E, Petrovas C. The Lymph Node in HIV Pathogenesis. Curr HIV/AIDS Rep. 2017;14(4):133–40.

8. Tas JM, Mesin L, Pasqual G, Targ S, Jacobsen JT, Mano YM, et al. Visualizing antibody affinity maturation in germinal centers. Science. 2016;351(6277):1048–54.

9. Victora GD, Nussenzweig MC. Germinal Centers. Annu Rev Immunol. 2022;40:413–42.

10. Padhan K, Moysi E, Noto A, Chassiakos A, Ghneim K, Perra MM, et al. Acquisition of optimal TFH cell function is defined by specific molecular, positional, and TCR dynamic signatures. Proc Natl Acad Sci U S A. 2021;118(18).

11. Podestà MA, Cavazzoni CB, Hanson BL, Bechu ED, Ralli G, Clement RL, et al. Stepwise differentiation of follicular helper T cells reveals distinct developmental and functional states. Nat Commun. 2023;14(1):7712.

12. Liu X, Yan X, Zhong B, Nurieva RI, Wang A, Wang X, et al. Bcl6 expression specifies the T follicular helper cell program in vivo. J Exp Med. 2012;209(10):1841–52, s1–24.

13. Song W, Craft J. T Follicular Helper Cell Heterogeneity. Annu Rev Immunol. 2024;42(1):127–52.

14. Georgakis S, Orfanakis M, Fenwick C, Brenna C, Burgermeister S, Lindsay H, et al. Delineation of the Human Germinal Centre Immune Landscape Using Multiplex Imaging Analysis. Immunology. 2025;176(1):87–104.

15. Moysi E, Sharma AA, O’Dell S, Georgakis S, Del Rio Estrada PM, Khader G, et al. A favorable follicular helper CD4 T cell programming characterizes neutralization activity in chronic HIV infection. J Clin Invest. 2026.

16. Vinuesa CG. HIV and T follicular helper cells: a dangerous relationship. J Clin Invest. 2012;122(9):3059–62.

17. Marina-Zárate E, Sutton HJ, Lopez PG, Altheide TK, Bick M, Burton I, et al. Highly functional and prolonged germinal center T follicular helper cell responses are associated with enhanced neutralizing antibody development. Immunity. 2025;58(12):3094–112.e7.

18. Locci M, Havenar-Daughton C, Landais E, Wu J, Kroenke MA, Arlehamn CL, et al. Human circulating PD-1+CXCR3-CXCR5+ memory Tfh cells are highly functional and correlate with broadly neutralizing HIV antibody responses. Immunity. 2013;39(4):758–69.

19. Shulman Z, Gitlin AD, Targ S, Jankovic M, Pasqual G, Nussenzweig MC, et al. T follicular helper cell dynamics in germinal centers. Science. 2013;341(6146):673–7.

20. Dufour C, Gantner P, Fromentin R, Chomont N. The multifaceted nature of HIV latency. J Clin Invest. 2020;130(7):3381–90.

21. Banga R, Perreau M. The multifaceted nature of HIV tissue reservoirs. Curr Opin HIV AIDS. 2024;19(3):116–23.

22. Spiegel H, Herbst H, Niedobitek G, Foss HD, Stein H. Follicular dendritic cells are a major reservoir for human immunodeficiency virus type 1 in lymphoid tissues facilitating infection of CD4+ T-helper cells. Am J Pathol. 1992;140(1):15–22.

23. Lindqvist M, van Lunzen J, Soghoian DZ, Kuhl BD, Ranasinghe S, Kranias G, et al. Expansion of HIV-specific T follicular helper cells in chronic HIV infection. J Clin Invest. 2012;122(9):3271–80.

24. Petrovas C, Yamamoto T, Gerner MY, Boswell KL, Wloka K, Smith EC, et al. CD4 T follicular helper cell dynamics during SIV infection. J Clin Invest. 2012;122(9):3281–94.

25. Moysi E, Pallikkuth S, De Armas LR, Gonzalez LE, Ambrozak D, George V, et al. Altered immune cell follicular dynamics in HIV infection following influenza vaccination. J Clin Invest. 2018;128(7):3171–85.

26. Pasternak AO, van Paassen PM, Verschoor YL, Vroom J, van Dort KA, Maurer I, et al. Long-term effect of temporary ART initiated during primary HIV-1 infection on viral persistence. Nat Commun. 2025;16(1):6989.

27. Pasternak AO, Grijsen ML, Wit FW, Bakker M, Jurriaans S, Prins JM, et al. Cell-associated HIV-1 RNA predicts viral rebound and disease progression after discontinuation of temporary early ART. JCI Insight. 2020;5(6).

28. Guerville F, Vialemaringe M, Cognet C, Duffau P, Lazaro E, Cazanave C, et al. Mechanisms of systemic low-grade inflammation in HIV patients on long-term suppressive antiretroviral therapy: the inflammasome hypothesis. Aids. 2023;37(7):1035–46.

29. Banga R, Procopio FA, Lana E, Gladkov GT, Roseto I, Parsons EM, et al. Lymph node dendritic cells harbor inducible replication-competent HIV despite years of suppressive ART. Cell Host Microbe. 2023;31(10):1714–31.e9.

30. Sun W, Gao C, Hartana CA, Osborn MR, Einkauf KB, Lian X, et al. Phenotypic signatures of immune selection in HIV-1 reservoir cells. Nature. 2023;614(7947):309–17.

31. Perreau M, Savoye AL, De Crignis E, Corpataux JM, Cubas R, Haddad EK, et al. Follicular helper T cells serve as the major CD4 T cell compartment for HIV-1 infection, replication, and production. J Exp Med. 2013;210(1):143–56.

32. Buggert M, Nguyen S, Salgado-Montes de Oca G, Bengsch B, Darko S, Ransier A, et al. Identification and characterization of HIV-specific resident memory CD8(+) T cells in human lymphoid tissue. Sci Immunol. 2018;3(24).

33. Lichterfeld M, Kaufmann DE, Yu XG, Mui SK, Addo MM, Johnston MN, et al. Loss of HIV-1-specific CD8+ T cell proliferation after acute HIV-1 infection and restoration by vaccine-induced HIV-1-specific CD4+ T cells. J Exp Med. 2004;200(6):701–12.

34. Petrovas C, Ferrando-Martinez S, Gerner MY, Casazza JP, Pegu A, Deleage C, et al. Follicular CD8 T cells accumulate in HIV infection and can kill infected cells in vitro via bispecific antibodies. Sci Transl Med. 2017;9(373).

35. Walker CM, Moody DJ, Stites DP, Levy JA. CD8+ lymphocytes can control HIV infection in vitro by suppressing virus replication. Science. 1986;234(4783):1563–6.

36. Collins DR, Hitschfel J, Urbach JM, Mylvaganam GH, Ly NL, Arshad U, et al. Cytolytic CD8(+) T cells infiltrate germinal centers to limit ongoing HIV replication in spontaneous controller lymph nodes. Sci Immunol. 2023;8(83):eade5872.

37. Quigley M, Pereyra F, Nilsson B, Porichis F, Fonseca C, Eichbaum Q, et al. Transcriptional analysis of HIV-specific CD8+ T cells shows that PD-1 inhibits T cell function by upregulating BATF. Nat Med. 2010;16(10):1147–51.

38. Collins DR, Urbach JM, Racenet ZJ, Arshad U, Power KA, Newman RM, et al. Functional impairment of HIV-specific CD8(+) T cells precedes aborted spontaneous control of viremia. Immunity. 2021;54(10):2372–84.e7.

39. Rutishauser RL, Deguit CDT, Hiatt J, Blaeschke F, Roth TL, Wang L, et al. TCF-1 regulates HIV-specific CD8+ T cell expansion capacity. JCI Insight. 2021;6(3).

40. Peluso MJ, Sandel DA, Deitchman AN, Kim SJ, Dalhuisen T, Tummala HP, et al. Correlates of HIV-1 control after combination immunotherapy. Nature. 2026;650(8100):187–95.

41. Kiani Z, Urbach JM, Wisner H, Olatotse MJ, Chang DY, Acklin JA, et al. CD8(+) T cell stemness precedes post-intervention control of HIV viraemia. Nature. 2026;650(8100):196–204.

42. Okoye AA, Duell DD, Fukazawa Y, Varco-Merth B, Marenco A, Behrens H, et al. CD8+ T cells fail to limit SIV reactivation following ART withdrawal until after viral amplification. J Clin Invest. 2021;131(8).

43. Pampena MB, Samer S, Viox EG, Nguyen K, Deleage C, Kuri-Cervantes L, et al. Therapeutic CD8(+) T cell tissue retention and immunomodulation during ART interruption fail to prevent SIV rebound. Proc Natl Acad Sci U S A. 2025;122(33):e2501037122.

44. Migueles SA, Weeks KA, Nou E, Berkley AM, Rood JE, Osborne CM, et al. Defective human immunodeficiency virus-specific CD8+ T-cell polyfunctionality, proliferation, and cytotoxicity are not restored by antiretroviral therapy. J Virol. 2009;83(22):11876–89.

45. Ollerton MT, Folkvord JM, La Mantia A, Parry DA, Meditz AL, McCarter MD, et al. Follicular regulatory T cells eliminate HIV-1-infected follicular helper T cells in an IL-2 concentration dependent manner. Front Immunol. 2022;13:878273.

46. Yero A, Shi T, Farnos O, Routy JP, Tremblay C, Durand M, et al. Dynamics and epigenetic signature of regulatory T-cells following antiretroviral therapy initiation in acute HIV infection. EBioMedicine. 2021;71:103570.

47. Miles B, Miller SM, Folkvord JM, Kimball A, Chamanian M, Meditz AL, et al. Follicular regulatory T cells impair follicular T helper cells in HIV and SIV infection. Nat Commun. 2015;6:8608.

48. Huber A, Baas FS, van der Ven A, Dos Santos JC. Innate Immune Cell Functions Contribute to Spontaneous HIV Control. Curr HIV/AIDS Rep. 2024;22(1):6.

49. Li SY, Yin LB, Ding HB, Liu M, Lv JN, Li JQ, et al. Altered lipid metabolites accelerate early dysfunction of T cells in HIV-infected rapid progressors by impairing mitochondrial function. Front Immunol. 2023;14:1106881.

50. Talla A, Azevedo J, Latif MB, Enriquez AB, Sanchez GP, Pelletier AN, et al. Innate antiviral and immune functions associated with the HIV reservoir decay after anti-PD-1 therapy. Nat Med. 2026;32(2):505–17.

51. Martin-Gayo E, Gao C, Chen HR, Ouyang Z, Kim D, Kolb KE, et al. Immunological Fingerprints of Controllers Developing Neutralizing HIV-1 Antibodies. Cell Rep. 2020;30(4):984– 96.e4.

52. Mitchell JL, Takata H, Muir R, Colby DJ, Kroon E, Crowell TA, et al. Plasmacytoid dendritic cells sense HIV replication before detectable viremia following treatment interruption. J Clin Invest. 2020;130(6):2845–58.

53. Zhu L, Ji J, Xiao J, Wang F, Yu J, Liu Y, et al. Interferons in HIV-1 infection: mechanisms, antiviral potentials, and therapeutic challenges. Front Immunol. 2025;16:1736658.

54. Jacquelin B, Mayau V, Targat B, Liovat AS, Kunkel D, Petitjean G, et al. Nonpathogenic SIV infection of African green monkeys induces a strong but rapidly controlled type I IFN response. J Clin Invest. 2009;119(12):3544–55.

55. Doyle T, Goujon C, Malim MH. HIV-1 and interferons: who’s interfering with whom? Nat Rev Microbiol. 2015;13(7):403–13.

56. Zhen A, Rezek V, Youn C, Lam B, Chang N, Rick J, et al. Targeting type I interferon-mediated activation restores immune function in chronic HIV infection. J Clin Invest. 2017;127(1):260–8.

57. Swainson LA, Sharma AA, Ghneim K, Ribeiro SP, Wilkinson P, Dunham RM, et al. IFN-α blockade during ART-treated SIV infection lowers tissue vDNA, rescues immune function, and improves overall health. JCI Insight. 2022;7(5).

58. Liu X, Zhang L, Li X, Chen L, Lu L, Yang Y, et al. Single-cell multi-omics profiling uncovers the immune heterogeneity in HIV-infected immunological non-responders. EBioMedicine. 2025;115:105667.

59. Pelletier AN, Sanchez GP, Izmirly A, Watson M, Di Pucchio T, Carvalho KI, et al. A pre-vaccination immune metabolic interplay determines the protective antibody response to a dengue virus vaccine. Cell Rep. 2024;43(7):114370.

60. Ramon S, Baker SF, Sahler JM, Kim N, Feldsott EA, Serhan CN, et al. The specialized proresolving mediator 17-HDHA enhances the antibody-mediated immune response against influenza virus: a new class of adjuvant? J Immunol. 2014;193(12):6031–40.

61. Schaeuble K, Cannelle H, Favre S, Huang HY, Oberle SG, Speiser DE, et al. Attenuation of chronic antiviral T-cell responses through constitutive COX2-dependent prostanoid synthesis by lymph node fibroblasts. PLoS Biol. 2019;17(7):e3000072.

62. Duffney PF, Falsetta ML, Rackow AR, Thatcher TH, Phipps RP, Sime PJ. Key roles for lipid mediators in the adaptive immune response. J Clin Invest. 2018;128(7):2724–31.

63. Ryan EP, Pollock SJ, Murant TI, Bernstein SH, Felgar RE, Phipps RP. Activated human B lymphocytes express cyclooxygenase-2 and cyclooxygenase inhibitors attenuate antibody production. J Immunol. 2005;174(5):2619–26.

64. Torres RM, Cyster J. Lipid mediators in the regulation of innate and adaptive immunity. Immunol Rev. 2023;317(1):4–7.

65. Kalinski P. Regulation of immune responses by prostaglandin E2. J Immunol. 2012;188(1):21–8.

66. Sreeramkumar V, Fresno M, Cuesta N. Prostaglandin E2 and T cells: friends or foes? Immunol Cell Biol. 2012;90(6):579–86.

67. Chen JH, Perry CJ, Tsui YC, Staron MM, Parish IA, Dominguez CX, et al. Prostaglandin E2 and programmed cell death 1 signaling coordinately impair CTL function and survival during chronic viral infection. Nat Med. 2015;21(4):327–34.

68. Hayes MM, Lane BR, King SR, Markovitz DM, Coffey MJ. Prostaglandin E(2) inhibits replication of HIV-1 in macrophages through activation of protein kinase A. Cell Immunol. 2002;215(1):61–71.

69. Coulombe F, Jaworska J, Verway M, Tzelepis F, Massoud A, Gillard J, et al. Targeted prostaglandin E2 inhibition enhances antiviral immunity through induction of type I interferon and apoptosis in macrophages. Immunity. 2014;40(4):554–68.

70. Clemente MI, Álvarez S, Serramía MJ, Martínez-Bonet M, Muñoz-Fernández M. Prostaglandin E2 reduces the release and infectivity of new cell-free virions and cell-to-cell HIV-1 transfer. PLoS One. 2014;9(2):e85230.

71. Prebensen C, Trøseid M, Ueland T, Dahm A, Sandset PM, Aaberge I, et al. Immune activation and HIV-specific T cell responses are modulated by a cyclooxygenase-2 inhibitor in untreated HIV-infected individuals: An exploratory clinical trial. PLoS One. 2017;12(5):e0176527.

72. Pettersen FO, Torheim EA, Dahm AE, Aaberge IS, Lind A, Holm M, et al. An exploratory trial of cyclooxygenase type 2 inhibitor in HIV-1 infection: downregulated immune activation and improved T cell-dependent vaccine responses. J Virol. 2011;85(13):6557–66.

73. Aandahl EM, Aukrust P, Skålhegg BS, Müller F, Frøland SS, Hansson V, et al. Protein kinase A type I antagonist restores immune responses of T cells from HIV-infected patients. Faseb j. 1998;12(10):855–62.

74. Sips M, Gerlo S, De Clercq L, Gomez EA, Colas RA, Dalli J, et al. Distinct immune profiles of HIV-infected subjects are linked to specific lipid mediator signature. Immun Inflamm Dis. 2022;10(6):e629.

75. Gerner MY, Kastenmuller W, Ifrim I, Kabat J, Germain RN. Histo-cytometry: a method for highly multiplex quantitative tissue imaging analysis applied to dendritic cell subset microanatomy in lymph nodes. Immunity. 2012;37(2):364–76.

76. Xia Y, Sandor K, Pai JA, Daniel B, Raju S, Wu R, et al. BCL6-dependent TCF-1(+) progenitor cells maintain effector and helper CD4(+) T cell responses to persistent antigen. Immunity. 2022;55(7):1200–15.e6.

77. Clouthier DL, Zhou AC, Wortzman ME, Luft O, Levy GA, Watts TH. GITR intrinsically sustains early type 1 and late follicular helper CD4 T cell accumulation to control a chronic viral infection. PLoS Pathog. 2015;11(1):e1004517.

78. Ming S, Yin H, Li X, Gong S, Zhang G, Wu Y. GITR Promotes the Polarization of TFH-Like Cells in Helicobacter pylori-Positive Gastritis. Front Immunol. 2021;12:736269.

79. Baumjohann D, Preite S, Reboldi A, Ronchi F, Ansel KM, Lanzavecchia A, et al. Persistent antigen and germinal center B cells sustain T follicular helper cell responses and phenotype. Immunity. 2013;38(3):596–605.

80. Zhu F, McMonigle RJ, Schroeder AR, Xia X, Figge D, Greer BD, et al. Spatiotemporal resolution of germinal center Tfh cell differentiation and divergence from central memory CD4(+) T cell fate. Nat Commun. 2023;14(1):3611.

81. Haberman AM, Gonzalez DG, Wong P, Zhang TT, Kerfoot SM. Germinal center B cell initiation, GC maturation, and the coevolution of its stromal cell niches. Immunol Rev. 2019;288(1):10–27.

82. He R, Hou S, Liu C, Zhang A, Bai Q, Han M, et al. Follicular CXCR5-expressing CD8(+) T cells curtail chronic viral infection. Nature. 2016;537(7620):412–28.

83. Mylvaganam GH, Rios D, Abdelaal HM, Iyer S, Tharp G, Mavigner M, et al. Dynamics of SIV-specific CXCR5+ CD8 T cells during chronic SIV infection. Proc Natl Acad Sci U S A. 2017;114(8):1976–81.

84. Chen Y, Yu M, Zheng Y, Fu G, Xin G, Zhu W, et al. CXCR5(+)PD-1(+) follicular helper CD8 T cells control B cell tolerance. Nat Commun. 2019;10(1):4415.

85. Miller SM, Miles B, Guo K, Folkvord J, Meditz AL, McCarter MD, et al. Follicular Regulatory T Cells Are Highly Permissive to R5-Tropic HIV-1. J Virol. 2017;91(17).

86. Macintyre AN, Gerriets VA, Nichols AG, Michalek RD, Rudolph MC, Deoliveira D, et al. The glucose transporter Glut1 is selectively essential for CD4 T cell activation and effector function. Cell Metab. 2014;20(1):61–72.

87. Dubois B, Barthélémy C, Durand I, Liu YJ, Caux C, Brière F. Toward a role of dendritic cells in the germinal center reaction: triggering of B cell proliferation and isotype switching. J Immunol. 1999;162(6):3428–36.

88. Bodelet J. Statistical quantile learning for large additive latent variable models. Journal of the American Statistical Association. 2025:1–22.

89. Wang H, Geng J, Wen X, Bi E, Kossenkov AV, Wolf AI, et al. The transcription factor Foxp1 is a critical negative regulator of the differentiation of follicular helper T cells. Nat Immunol. 2014;15(7):667–75.

90. Zou D, Li XC, Chen W. Beyond T-cell subsets: stemness and adaptation redefining immunity and immunotherapy. Cell Mol Immunol. 2025;22(9):957–74.

91. Wang F, Cheng F, Zheng F. Stem cell like memory T cells: A new paradigm in cancer immunotherapy. Clin Immunol. 2022;241:109078.

92. Teng CF, Jeng LB, Shyu WC. Role of Insulin-like Growth Factor 1 Receptor Signaling in Stem Cell Stemness and Therapeutic Efficacy. Cell Transplant. 2018;27(9):1313–9.

93. Locci M, Wu JE, Arumemi F, Mikulski Z, Dahlberg C, Miller AT, et al. Activin A programs the differentiation of human TFH cells. Nat Immunol. 2016;17(8):976–84.

94. Skartsis N, Peng Y, Ferreira LMR, Nguyen V, Ronin E, Muller YD, et al. IL-6 and TNFα Drive Extensive Proliferation of Human Tregs Without Compromising Their Lineage Stability or Function. Front Immunol. 2021;12:783282.

95. Legler DF, Bruckner M, Uetz-von Allmen E, Krause P. Prostaglandin E2 at new glance: novel insights in functional diversity offer therapeutic chances. Int J Biochem Cell Biol. 2010;42(2):198– 201.

96. Park JY, Pillinger MH, Abramson SB. Prostaglandin E2 synthesis and secretion: the role of PGE2 synthases. Clin Immunol. 2006;119(3):229–40.

97. Battivelli E, Dahabieh MS, Abdel-Mohsen M, Svensson JP, Tojal Da Silva I, Cohn LB, et al. Distinct chromatin functional states correlate with HIV latency reactivation in infected primary CD4(+) T cells. Elife. 2018;7.

98. Georgakis S, Orfanakis M, Brenna C, Burgermeister S, Del Rio Estrada PM, González-Navarro M, et al. Follicular Immune Landscaping Reveals a Distinct Profile of FOXP3(hi)CD4(hi) T Cells in Treated Compared to Untreated HIV. Vaccines (Basel). 2024;12(8).

99. McNally A, Hill GR, Sparwasser T, Thomas R, Steptoe RJ. CD4+CD25+ regulatory T cells control CD8+ T-cell effector differentiation by modulating IL-2 homeostasis. Proc Natl Acad Sci U S A. 2011;108(18):7529–34.

100. Chung Y, Tanaka S, Chu F, Nurieva RI, Martinez GJ, Rawal S, et al. Follicular regulatory T cells expressing Foxp3 and Bcl-6 suppress germinal center reactions. Nat Med. 2011;17(8):983–8.

101. Yu M, Qu M, Wang Z, Zhen C, Yang B, Zhang Y, et al. Dysfunction and Metabolic Reprogramming of Gut Regulatory T Cells in HIV-Infected Immunological Non-Responders. Cells. 2025;14(15).

102. Lu J, Liang Y, Meng H, Zhang A, Zhao J, Zhang C. Metabolic Controls on Epigenetic Reprogramming in Regulatory T Cells. Front Immunol. 2021;12:728783.

103. de Paula HHS, Ferreira ACG, Caetano DG, Delatorre E, Teixeira SLM, Coelho LE, et al. Reduction of inflammation and T cell activation after 6 months of cART initiation during acute, but not in early chronic HIV-1 infection. Retrovirology. 2018;15(1):76.

104. Anzinger JJ, Butterfield TR, Angelovich TA, Crowe SM, Palmer CS. Monocytes as regulators of inflammation and HIV-related comorbidities during cART. J Immunol Res. 2014;2014:569819.

105. Grootveld AK, Kyaw W, Panova V, Lau AWY, Ashwin E, Seuzaret G, et al. Apoptotic cell fragments locally activate tingible body macrophages in the germinal center. Cell. 2023;186(6):1144– 61.e18.

106. Gurwicz N, Stoler-Barak L, Schwan N, Bandyopadhyay A, Meyer-Hermann M, Shulman Z. Tingible body macrophages arise from lymph node-resident precursors and uptake B cells by dendrites. J Exp Med. 2023;220(4).

107. Liu T, Yang Q, Cao YJ, Yuan WM, Lei AH, Zhou P, et al. Cyclooxygenase-1 Regulates the Development of Follicular Th Cells via Prostaglandin E(2). J Immunol. 2019;203(4):864–72.

108. Mahic M, Yaqub S, Johansson CC, Taskén K, Aandahl EM. FOXP3+CD4+CD25+ adaptive regulatory T cells express cyclooxygenase-2 and suppress effector T cells by a prostaglandin E2-dependent mechanism. J Immunol. 2006;177(1):246–54.

109. Bekeredjian-Ding I, Schäfer M, Hartmann E, Pries R, Parcina M, Schneider P, et al. Tumour-derived prostaglandin E and transforming growth factor-beta synergize to inhibit plasmacytoid dendritic cell-derived interferon-alpha. Immunology. 2009;128(3):439–50.

110. Baratelli F, Lin Y, Zhu L, Yang SC, Heuzé-Vourc’h N, Zeng G, et al. Prostaglandin E2 induces FOXP3 gene expression and T regulatory cell function in human CD4+ T cells. J Immunol. 2005;175(3):1483–90.

111. Boniface K, Bak-Jensen KS, Li Y, Blumenschein WM, McGeachy MJ, McClanahan TK, et al. Prostaglandin E2 regulates Th17 cell differentiation and function through cyclic AMP and EP2/EP4

112. Betz M, Fox BS. Prostaglandin E2 inhibits production of Th1 lymphokines but not of Th2 lymphokines. J Immunol. 1991;146(1):108–13.

113. Boraschi D, Censini S, Bartalini M, Scapigliati G, Barbarulli G, Vicenzi E, et al. Interferon inhibits prostaglandin biosynthesis in macrophages: effects on arachidonic acid metabolism. J Immunol. 1984;132(4):1987–92.

114. Wahl LM, Corcoran ME, Mergenhagen SE, Finbloom DS. Inhibition of phospholipase activity in human monocytes by IFN-gamma blocks endogenous prostaglandin E2-dependent collagenase production. J Immunol. 1990;144(9):3518–22.

115. Hannigan GE, Williams BR. Signal transduction by interferon-alpha through arachidonic acid metabolism. Science. 1991;251(4990):204–7.

116. Hong DS, Parikh A, Shapiro GI, Varga A, Naing A, Meric-Bernstam F, et al. First-in-human phase I study of immunomodulatory E7046, an antagonist of PGE(2)-receptor E-type 4 (EP4), in patients with advanced cancers. J Immunother Cancer. 2020;8(1).

117. Pietrantonio F, Morano F, Niger M, Ghelardi F, Chiodoni C, Palazzo M, et al. The Prostaglandin EP4 Antagonist Vorbipiprant Combined with PD-1 Blockade for Refractory Microsatellite-Stable Metastatic Colorectal Cancer: A Phase Ib/IIa Trial. Clin Cancer Res. 2025;31(4):649–58.

118. Kvale D, Ormaasen V, Kran AM, Johansson CC, Aukrust P, Aandahl EM, et al. Immune modulatory effects of cyclooxygenase type 2 inhibitors in HIV patients on combination antiretroviral treatment. Aids. 2006;20(6):813–20.

119. Buzon MJ, Sun H, Li C, Shaw A, Seiss K, Ouyang Z, et al. HIV-1 persistence in CD4+ T cells with stem cell-like properties. Nat Med. 2014;20(2):139–42.

120. Kunzli M, Schreiner D, Pereboom TC, Swarnalekha N, Litzler LC, Lotscher J, et al. Long-lived T follicular helper cells retain plasticity and help sustain humoral immunity. Sci Immunol. 2020;5(45).

121. Sandler NG, Bosinger SE, Estes JD, Zhu RT, Tharp GK, Boritz E, et al. Type I interferon responses in rhesus macaques prevent SIV infection and slow disease progression. Nature. 2014;511(7511):601–5.

122. Wu VH, Yung BS, Faraji F, Saddawi-Konefka R, Wang Z, Wenzel AT, et al. The GPCR-Galpha(s)-PKA signaling axis promotes T cell dysfunction and cancer immunotherapy failure. Nat Immunol. 2023;24(8):1318–30.

123. Mallarino-Haeger C, Pino M, Viox EG, Pagliuzza A, King CT, Nguyen K, et al. HIV-1 DNA and Immune Activation Levels Differ for Long-Lived T-Cells in Lymph Nodes, Compared with Peripheral Blood, during Antiretroviral Therapy. J Virol. 2023;97(4):e0167022.

124. Georgakis S, Ioannidou K, Mora BB, Orfanakis M, Brenna C, Muller YD, et al. Cellular and molecular determinants mediating the dysregulated germinal center immune dynamics in systemic lupus erythematosus. Front Immunol. 2025;16:1530327.

125. Law CW, Chen Y, Shi W, Smyth GK. voom: Precision weights unlock linear model analysis tools for RNA-seq read counts. Genome Biol. 2014;15(2):R29.

126. Wu D, Smyth GK. Camera: a competitive gene set test accounting for inter-gene correlation. Nucleic Acids Res. 2012;40(17):e133.

127. Delporte M, Lambrechts L, Blomme EE, van Snippenberg W, Rutsaert S, Verschoore M, et al. Integrative Assessment of Total and Intact HIV-1 Reservoir by a 5-Region Multiplexed Rainbow DNA Digital PCR Assay. Clin Chem. 2025;71(1):203–14.

128. Radtke AJ, Kandov E, Lowekamp B, Speranza E, Chu CJ, Gola A, et al. IBEX: A versatile multiplex optical imaging approach for deep phenotyping and spatial analysis of cells in complex tissues. Proc Natl Acad Sci U S A. 2020;117(52):33455–65.

129. Deleage C, Wietgrefe SW, Del Prete G, Morcock DR, Hao XP, Piatak M, Jr., et al. Defining HIV and SIV Reservoirs in Lymphoid Tissues. Pathog Immun. 2016;1(1):68–106.

130. Garcia JV, Miller AD. Serine phosphorylation-independent downregulation of cell-surface CD4 by nef. Nature. 1991;350(6318):508–11.

131. Battivelli E, Dahabieh MS, Abdel-Mohsen M, Svensson JP, Tojal Da Silva I, Cohn LB, et al. Distinct chromatin functional states correlate with HIV latency reactivation in infected primary CD4(+) T cells. Elife. 2018;7.

132. Battivelli E, Verdin E. HIV(GKO): A Tool to Assess HIV-1 Latency Reversal Agents in Human Primary CD4(+) T Cells. Bio Protoc. 2018;8(20).

133. Barde I, Salmon P, Trono D. Production and titration of lentiviral vectors. Curr Protoc Neurosci. 2010;Chapter 4:Unit 4.21.

