## Extended data for "An eicosanoid-enriched follicular microenvironment shapes germinal centre immune dynamics in antiretroviral-treated PLWH"

**Methods**

***Blood HIV DNA measurements***

Starting from 10-20x10^6^ cryopreserved PBMCs, CD4+ T-cells were enriched by negative selection using the EasySep Human CD4+ T-cell isolation kit (Stemcell Technologies, Vancouver, Canada). Genomic DNA (gDNA) was extracted using the QiaAmp DNA mini kit on the Qiacube (Qiagen, Hilden, Germany) with two elution steps of 50µL. DNA concentrations were determined using 2 µL of the eluted DNA with the Qubit 2.0 fluorometer using the Qubit™ 1X dsDNA Broad Range (BR) Assay kit (Thermofisher Scientific, USA). Samples were stored at -20°C prior to dPCR quantification.

HIV DNA levels were quantified in triplicate by dPCR using the Rainbow proviral HIV-1 DNA assay on the QIAcuity Four platform (Qiagen, Hilden, Germany) (127). Depending on concentration, genomic DNA (gDNA) inputs ranged from 384 to 1022 ng per replicate, with elution volumes between 12 and 19 µL. To ensure accurate quantification, HIV-1 DNA levels were normalized against the reference gene RPP30, measured in duplicate using 5 µL of a 1/100 dilution for each sample. Total HIV DNA levels were assessed by the RU5 region in the Rainbow assay. Automatic thresholds were calculated with the Rainbow Shiny tool and adapted if needed per sample when a threshold intersected a cluster of positive partitions. Final values were reported as copies per million CD4+ T-cells.

***mIF data acquisition***

Images were acquired using a Leica Stellaris 8 (SP8) confocal system, equipped with Leica Application Suite X (LAS-X/4.6.1.27508) software, at 512 × 512-pixel density, 0.75× optical zoom and a z-step of 1 um using a 20× objective (NA). Frame averaging or summing was never used while obtaining the images. At least 90% of each section was imaged, to ensure an accurate representation and minimize selection bias. Tissues stained with a single antibody fluorophore combination were used to create a compensation matrix via the Leica LAS-AF Channel Dye Separation module (Leica Microsystems), which was used to fluorophore spillover correction (when present), in accordance with the user manual. If the results of the dye separation were not optimal, the LAS-AF Channel Dye Separation module was used manually.

***Image Alignment and Registration- Cell segmentation***

The alignment of the images generated during the two different imaging cycles was performed using SimpleITK (128) as an Imaris extension (Imaris software version 9.9.0, Biplane). To facilitate registration, we utilized one or two common channels present in both imaging cycles (SYTO or CD57-BV421). After successful alignment, the Surface Creation module of Imaris was used to generate 3-dimensional segmented surfaces (based on the nuclear signal intensity) of spillover-corrected images. The segmented cells were then processed using the Imaris filtering module with different combinations of filtering types, based on the mean and median intensities of the channels to exclude artifacts characterized by uniform staining across the segmented area. Areas with uniform/non-specific staining were excluded among the different tissues. Data generated, such as average voxel intensities for all antibody-related channels, in addition to the volume and sphericity of the 3-dimensional surfaces, were exported in Microsoft Excel format.

***HIV DNAscope***

DNAscope in situ hybridization (ISH) for HIV DNA was performed using the HIV Clade B sense DNA probe (Cat. No. 425531, ACD) and the RNAscope Red kit (Cat. No. 322360, ACD), with minor modifications based on the protocol described in (129). FFPE sections were deparaffinized using the standard Ventana Discovery protocol, followed by antigen retrieval (DAKO; 15 min, 100°C), Protease III treatment (20 min, 40°C), and incubation with 3% H₂O₂ at room temperature. Sections were hybridized with the HIV probe for 4 h at 40°C and processed through the RNAscope amplification steps. After a 30-min blocking step, DNAscope-compatible protein markers (CD3 [**Extended Data Table 3**], CD20 [**Extended Data Table 3**] and CD14 [CatNo 114R-14] were diluted in Antibody Diluent/Blocker (90 min, room temperature). Warp Red Chromogen kit was then applied to visualize HIV DNA, followed by DAPI counterstaining and mounting with DAKO mounting medium. The protocol was validated using the latently HIV-infected ACH-2 cell line (data not shown).

Images were acquired on a Leica Stellaris SP8 confocal microscope (LAS X v4.6.1) using a 40×/0.75 NA objective (512 × 512 pixels, 0.75× optical zoom, 0.8-μm z-step, no frame averaging). At least 80% of each tissue section was imaged to minimize sampling bias. Because Fast Red is not a true fluorophore, its signal was detected using the ATTO 550 emission settings (580 ± 20 nm). Corresponding brightfield images of the same regions were acquired with a Leica THUNDER Imager to facilitate discrimination of true HIV-DNA signals from background.

Confocal and brightfield images were manually co-registered in Imaris v9.9.1. HIV-DNA-positive events were segmented in three dimensions using the Surface Creation module, while a colocalization channel generated from the nuclear and CD14 signals aided background identification. Sequential filtering based on object size, HIV-DNA intensity, colocalization, and visual comparison with the THUNDER brightfield images was applied to remove false-positive events. Finally, only HIV-DNA signals localized within CD3^hi^ cells were retained for our analysis. The validated HIV-DNA surfaces were then masked with the nuclear channel to generate the final filtered HIV-DNA signal for downstream analyses.

***Quantitative Imaging Analysis (Histocytometry)***

The excel files obtained from cell segmentation were converted to comma separated value (.CVS) files, and data were imported into FlowJo (version 10) for further analysis. Well-defined areas devoid of background staining were included in the analysis, and the data were quantified either as relative frequencies (%) or as cell densities (cell counts normalized to the imaged follicular and/or extrafollicular area). Imaging data were analysed either as individual samples or by applying a batch analysis. For the analysis of individual samples, manual hand gating was used for the identification of relevant cell subsets and the intensities of individual biomarkers used in gated populations were presented as histo-dotplots. Follicular areas were identified based on the density of CD20^hi/dim^, a biomarker specific for B cells, events. The cut-off values for the identification of cells expressing ‘high’ profile for a given biomarker (e.g. CD20, CD3, PD1) was determined based on the 2D plot distribution profile for this biomarker and the inspection of its intensity in the raw mIF image. The CD4 marker was included in our imaging panel initially, but it was excluded from our analysis because of its downregulation in PLWH (130). Histocytometry-analysed cells of interest were exported and imported into Imaris raw mIF image as segmented spots for the comparison/validation of these cells to their original counterparts. For batching unsupervised analysis, the same number (ID) was assigned to all segmented cells from a given tissue and all raw intensities exported from Imaris were normalized. Data normalization was performed using the StandardScaler from the sklearn library from Python 3.9.13. The normalized data was then reconstructed into a DataFrame, ensuring that excluded columns were preserved in their original form. Regions of Interest (ROIs-follicular areas) were identified for each donor and relevant cell subsets (e.g. follicular CD3^hi^PD1^hi^CD8^lo^, CD3^hi^CD8^hi^ etc) were manually gated. The same number of cells (400 cells for T_FH_) were concatenated together with the associated biomarkers under investigation. The concatenated ‘sample’ was processed using FlowJo10 modules/plugins (tSNE, FlowSOM (3.0.18) and Cluster Explorer) and the generated subpopulations were applied to all individual tissues included in the concatenated sample and were able to be identified based on the assigned ID number. FlowSOM analysis was carried out using SOM grid size 10x10, minimum spanning tree, a node scale of 100% and number of meta clusters 6. Heatmaps of generated subpopulations were generated using Python 3.9.13 to visualize the normalized data. The heatmap displayed a colour intensity range corresponding to markers expression levels.

***HIV DNA – Visium HD co-localization analysis***

HIV DNAscope images and Visium HD proximal sections from the same FFPE blocks of three cART-treated donors (cART-4, cART-6, cART-9) were analysed with a reproducible Snakemake workflow (Snakemake 9.6.2, pixi-locked environment).

*HIV DNA event detection and registration to Visium space.* Segmented DNAscope images were sum-projected across z and binarised (pixel value > 0); individual HIV DNA events were defined as 8-connected components (scipy.ndimage.label) and reduced to their pixel centroids. DNAscope and Visium coordinate spaces were aligned by landmark-based affine registration: corresponding landmark pairs were placed manually on the DNAscope and Visium images in napari, and a least-squares affine transform was estimated per sample (skimage.transform.AffineTransform). Event centroids were mapped into Visium space with this transform. Because a single physical event can map onto several adjacent Visium HD bins, transformed events were deduplicated by complete-linkage hierarchical clustering with a 1 µm distance threshold, keeping one centroid per cluster.

*Visium HD data and follicle assignment.* Space Ranger v4.0 segmented outputs (cell segmentations, filtered cell-by-gene matrix, high-resolution tissue image) were loaded with spatialdata-io 0.7.0 (SpatialData 0.7.3), and all coordinates were expressed in the downscaled high-resolution pixel space; pixel size in micrometres was derived from the Space Ranger scale factors as microns_per_pixel / tissue_hires_scalef. Follicles were annotated manually as barcode lists; each segmented cell was labelled inside or outside a follicle accordingly, and every HIV DNA event inherited the label of its nearest segmented cell. All downstream analyses were run separately for events inside and outside follicles.

*Neighbourhood pseudobulk and gene-set scoring.* For each HIV DNA event, raw counts of all segmented cells with a centroid within a 50 µm radius were summed into a neighbourhood pseudobulk profile. The same neighbourhood pooling was applied to every segmented cell of the region, yielding a reference population of pseudobulk profiles. To restrict scoring to reliably measured genes, genes detected in ≥ 10 cells in any sample/region were retained (union across samples), and for each gene set a panel of its detected genes plus 2,000 randomly drawn control genes (seed 42) not belonging to that set was used. Observed and reference profiles were concatenated, count-normalised (10,000 counts per profile), log1p-transformed and scored jointly with scanpy.tl.score genes (scanpy 1.12.1), so that control-gene bins are identical for observed and reference profiles. The test statistic was the mean gene-set score across HIV DNA neighbourhoods.

*Permutation test.* Spatial enrichment was assessed using a permutation test with 1,000 permutations (seed 42). For each sample and region, the number of cells sampled without replacement from the reference population was matched to the number of observed HIV DNA events, and the mean pseudobulk score was calculated for each permutation. Two-sided empirical p-values were computed as 2 × min (P (null ≥ observed), P (null ≤ observed)) with a +1 correction. For display, the observed statistic is expressed as a normalised permutation rank, rank = 2 × mean (null < observed) − 1, ranging from −1 (below all permutations) to +1 (above all permutations); ranks are plotted per donor against the number of HIV DNA events in that region, separately for publicly available and curated gene pathways.

*Spatial overview figures.* For each donor, the Space Ranger high-resolution tissue image and cell segmentation outlines were rendered with spatialdata-plot 0.3.3; follicles are drawn as the convex hull of the centroids of their annotated cells, deduplicated HIV DNA events as points, and the 50 µm cell neighbourhood as a circle around each event.

***Cell Culture***

Cryopreserved tonsillar cell suspensions were thawed carefully in warm culture medium (RPMI-1640/l-glutamine [cat No 21875-034; Gibco], supplemented with 10% v/v heat-inactivated FBS, 100 IU/mL penicillin, 100 μg/mL streptomycin, and 10 mm HEPES [Gibco]), seeded in 12-well plates (8x10^5^/well) and cultured in a humidified 37°C 5% CO2 incubator for 20 hours. For T cell activation, well plates were coated with aCD3 (1ug/ml) overnight and aCD28(1ug/ml) was added after cell seeding. CpG-B (0.1μM, cat No tlrl-2006, Invivogen) was used for B cell activation. Wherever needed, PGE₂ (5 μM, cat No. 233560100, Thermo Scientific) diluted in DMSO was used. Cells treated with plain DMSO were used as mock control.

***Preparation of high-titer purified HIV-GKO virus stocks***

The dual reporter (EGFP and mKO2) HIVGKO vector developed by Battivelli et al (131) was obtained from Addgene. Viral particles were produced using an HIV CCR5-tropic envelope as previously described (131, 132). Briefly, viral particles were prepared by transfection of HEK 293T cells. For this purpose, 10-12 x10^6^ HEK 293T cells were seeded per 15cm tissue culture dish in DMEM media with 10% v/v FBS and Penicillin-Streptomicin, Gentamicin (50 μg/ml, GIBCO). The next day, cells were transfected with 70 μg of plasmid DNA, through Calcium phosphate transfection (Takara) as previously described (133). The next day, media was replaced with DMEM with 10% FCS. After 2 days, media was filtered at 0.45 uM, collected into 38.5mL ultraclear centrifuge tubes (Herolab 253050), centrifuged at 50,000 g for 2 hours at 16°C. Virus pellet was re-suspended in 1 mL PBS, aliquoted and snap frozen in dry ice and then at -80°C for later use. Viral titers were determined by the assessment of the percentage of EGFP+ activated CD4 T cells with increasing virus doses at day 4 post-transduction by flow cytometry. Endotoxin levels were below the detection limit (Limulus amebocyte lysate assay; Sigma).

***In-vitro* infection assays**

CD4 T cells from HIV uninfected individuals (N=4) were negatively selected (EasySep™ Human CD4^+^ T Cell Isolation Kit) from fresh buffy coats. 200,000 CD4 T cells were exposed or not to anti-CD3/CD28 mAbs (10ug/mL each) or to the combination of anti-CD3/CD28 mAbs (10ug/mL each) and PGE_2_ (0.5 uM) for 24 hours in 96 well-flat bottom plates. After 24 hours, CD4 T cells were infected with HIVGKO. Cultures were re-supplemented with PGE_2_ (0.5 uM). Finally, cells were washed with PBS, stained for live cells, fixed and assessed for productive (EGFP+) and latent infection (mKO2+) on day 4 post-transduction by flow cytometry.

***Flow cytometry***

For the ex-vivo experiments using cell suspensions from PLWH**,** cryopreserved lymph node mononuclear cells (LNMCs) were rapidly thawed by direct transfer into pre-warmed RPMI 1640 medium supplemented with 10% v/v fetal bovine serum (FBS). Cells were centrifuged (1,200 rpm, 6 min) and counted by trypan blue exclusion using an automated cell counter (Countess, Thermo Fisher Scientific). Cell viability and recovery were recorded for each donor. Cells were treated with Benzonase for downstream flow cytometry analyses. Cells were pelleted by centrifugation (1,600 rpm, 7 min, room temperature), resuspended in RPMI 1640 supplemented with 10% FBS v/v containing Benzonase (1 U ml⁻¹), and incubated for 15 min at 37 °C. Cells were subsequently washed with RPMI + 10% FBS, centrifuged, and resuspended in complete culture medium consisting of AIM-V supplemented with penicillin–streptomycin (1% v/v), HEPES (1% v/v), sodium pyruvate (1% v/v), non-essential amino acids (1% v/v), and 10% v/v serum replacement (Corning NU-IV). After Benzonase treatment, 1 × 10⁶ cells per donor were allocated for flow cytometry. Cells were centrifuged, resuspended in culture medium, and plated into V-bottom plates. Surface staining was performed for 20 min at room temperature. Samples were then fixed and permeabilized using the eBioscience™ Foxp3 / Transcription Factor Staining Buffer Set (Invitrogen, Cat. #00-5523-00) for 45 min at 4 °C, followed by intracellular staining for 45 min at 4 °C. Data were acquired on a BD FACSymphony A5 flow cytometer (BD Biosciences) using BD FACSDiva software.

For *in-vitro* experiments using tonsillar cryopreserved samples, cells were collected from cultures, washed with staining buffer and stained with anti‐human Abs (5x10^6^ cells per test) diluted in PBS-FBS 2% v/v-1%FcR v/v blocking reagent staining buffer. Surface staining was performed at 4 °C for 30 min. Cells were then washed with staining buffer and resuspended in 250 μL for acquisition. Acquisition was performed on an LSR II flow cytometer (BD Biosciences) driven by BD FACSDiva software. Acquired data were analysed using FlowJo v.10.9.0. LIVE/DEAD Fixable Aqua Dead Cell Stain was used to exclude dead cells. Cells from both *ex-vivo* and *in-vitro* cells were phenotyped using the panels in **Extended Data Table 4**. Cytokines in matched culture supernatants were measured using the Luminex ProcartaPlex immunoassay as part of the routine diagnostic workflow of the CHUV Immunology Laboratory.

***scRNA data processing, clustering, cell type annotation and analysis***

Gene expression counts of the eight lymph node samples (4 cART, 4 Vir) were analysed in Python with Scanpy (v1.11.4) and AnnData (v0.12.3). Cells with >10% mitochondrial reads were removed, and putative doublets were identified with Scrublet (v0.2.3; expected doublet rate 0.06, min_counts = 2, min_cells = 3, min_gene_variability_pctl = 85, 30 principal components) and excluded. Cells expressing fewer than 400 genes and genes detected in fewer than 20 cells were discarded, leaving 42,936 nuclei for downstream analysis. Counts were normalised to 10,000 counts per cell, log1p-transformed, and the 3,000 most highly variable genes were used for principal component analysis (30 PCs, ARPACK solver). A k-nearest-neighbour graph (k = 15, 30 PCs) was built, embedded in two dimensions with UMAP, and partitioned with the Leiden algorithm (igraph implementation, python-igraph v0.11.9) at a coarse resolution of 0.08 to resolve major lineages. All steps used a fixed random seed (42) to ensure reproducible embeddings and clusterings.

Cluster identities were assigned manually. For every cluster, differentially expressed genes were computed against all remaining cells with a Wilcoxon rank-sum test (sc.tl.rank_genes_groups), and the top-ranked genes were inspected as dot plots (expression scaled per gene) together with curated marker panels projected onto the UMAP. To reduce the impact of sparse count noise in the marker UMAPs, expression values were binned into 50 percentile ranks prior to plotting. At the coarse resolution this defined four major populations: T cells (n = 23,668; *CD3D, CD3E, TRAC, THEMIS, BCL11B*), B cells (n = 17,622; *MS4A1, CD19, PAX5, BANK1, EBF1*), monocytes/macrophages/dendritic cells (n = 1,176; *CD14, CST3, TYMP, ITGAX*) and plasmacytoid dendritic cells (n = 470; *TCF4, IRF8, RUNX2, TLR7*).

T cells and B cells were then subsetted and re-analysed independently using the same normalisation, feature-selection, PCA, neighbourhood and UMAP parameters as above, so that each lineage is displayed on its own embedding computed from lineage-specific highly variable genes. Leiden clustering was performed over a range of resolutions (0.2–1.0) and the resolution giving stable, marker-supported clusters was retained (0.4 for T cells, nine clusters; 0.2 for B cells, seven clusters). T cell clusters were annotated as CD4 T cells (*CD4, TCF7, LEF1, IL7R*; n = 12,334), CD8 T cells (*CD8A, CD8B, CCL5, GZMK, PRF1, GZMB*; n = 7,386), regulatory T cells (*FOXP3, IL2RA, IKZF2, CTLA4, TNFRSF18*; n = 1,334), follicular helper T cells (*CXCR5, PDCD1, BCL6, ICOS, TOX2*; n = 1,212), proliferating T cells (*MKI67, PCNA, TOP2A, TYMS*; n = 996) and NK cells (*GNLY, NKG7, KLRD1, NCAM1*, low CD3D/CD3E; n = 406). B cell clusters were annotated as naive B cells (*IGHD, IGHM, FCER2, CR2*; n = 11,000), dark zone germinal centre B cells (*AICDA, CXCR4, MKI67, PCNA, FOXO1, BCL6*; n = 3,148), light zone germinal centre B cells (*CD83, CD40, ICAM1, ICOSLG, IL21R*; n = 2,358), tissue memory/age-associated B cells (*FCRL4, FCRL5, TNFRSF13B, ITGAX, TBX21, ZEB2*; n = 611) and plasma cells (*MZB1, XBP1*, *PRDM1, IRF4, IGHG1*; n = 505). The lineage-level labels were finally merged into a single per-cell annotation, in which T and B cells carry their fine-grained subtype label, and this annotation was used for all downstream analyses.

We tested for differential gene expression between cART and Vir samples using edgeR with counts pseudobulked per donor and cell type. Then, we tested gene set enrichment using limma fry. We observed that the p-value distributions of the fry results differed between annotated cell types. Therefore, to compare the relative importance of gene set differences between cART and viremic donors across cell types, we show gene set ranks rather than p-values for gene sets that were significant (Benjamini-Hochberg false discovery rate <= 0.05). For the comparison with public data, we downloaded the processed data from the GEO (GSE288212). The authors provided us with their cell type annotations, which we used in differential and gene set enrichment analyses as described for the sc-RNAseq.

| **ID** | **Age** | **Tissue Location** | **Sex** |
| --- | --- | --- | --- |
| HIVneg1 | 71 | Axillary | M |
| HIVneg2 | 49 | Inguinal | M |
| HIVneg3 | 25 | Inguinal | F |
| HIVneg4 | 78 | Inguinal | M |

**Extended Data Table 1**

| **ID** | **Age** | **Tissue Location** | **Sex** | **CD4 counts**  **(cells/uL)** | **CD8 counts**  **(cells/uL)** | **Treatment Duration** | **LogpVL** |
| --- | --- | --- | --- | --- | --- | --- | --- |
| Vir-1 | 28 | Cervical | M | 777 | 2195 | No treatment | 4.49 |
| Vir-2 | 27 | Cervical | M | 719 | 1831 | No treatment | 4.69 |
| Vir-3 | 35 | Cervical | M | 491 | 2284 | No treatment | 4.49 |
| Vir-4 | 31 | Inguinal | M | 281 | 2142 | No treatment | 6.29 |
| Vir-5 | 33 | Cervical | M | 499 | 2211 | No treatment | 6.06 |
| Vir-6 | 24 | Inguinal | M | 453 | 3623 | No treatment | 6.84 |
| Vir-7 | 37 | Inguinal | M | 391 | 1158 | No treatment | 6.25 |
| Vir-8 | 22 | Cervical | M | 476 | 465 | No treatment | 5.75 |
| Vir-9 | 29 | Inguinal | M | 545 | 830 | No treatment | 5.41 |
| Vir-10 | 20 | Cervical | M | 290 | 881 | No treatment | 7 |
| Vir-11 | 27 | Cervical | M | 520 | 175 | No treatment | 5.26 |
| Vir-12 | 29 | Cervical | M | 347 | 756 | No treatment | 7 |

| **ID** | **Age** | **Tissue Location** | **Sex** | **CD4 counts**  **(cells/uL)** | **HIV RNA copies/mL** | **ART Duration** | **Current ART Regimen** | **Years HIV+** | **HIV DNA**  **cp/10^6^ CD4 cells** |
| --- | --- | --- | --- | --- | --- | --- | --- | --- | --- |
| cART-1 | 58 | Inguinal | M | 636 | ND | 15 | DRV/RTV/FTC/TDF | 19 | 564.686 |
| cART-2 | 57 | Inguinal | M | 500 | <40 | 17 | DRV/RTV/ABC/FTC/TDF | 29 | 593.426 |
| cART-3 | 53 | Inguinal | M | 441 | ND | 10 | DTG/FTC/TDF | 22 |  |
| cART-4 | 37 | Inguinal | M | 662 | ND | 4 | EFV/FTC/TDF | 5 | 242.679 |
| cART-5 | 51 | Inguinal | M | 805 | ND | 10 | DTG/FTC/TDF | 17 |  |
| cART-6 | 33 | Inguinal | M | 1121 | ND | 2y 1mo | EVG/COBI/FTC/TDF | 3y5mo | 3296.14 |
| cART-7 | 44 | Inguinal | M | 545 | ND | 7 | ATV/RTV/FTC/TDF | 16 |  |
| cART-8 | 46 | Inguinal | M | 493 | ND | 3 | EVT/COBI/3TC/TDF | 3 | 615.037 |
| cART-9 | 63 | Inguinal | M | 276 | <40 | 4 | RPV/DRV/RTV/DTG | 14 | 3183.02 |
| cART-10 | 59 | Inguinal | M | 587 | ND | 10 | Atripla | 10 |  |
| cART-11 | 53 | Inguinal | F | 454 | ND | 11 | Triumeq | 26 |  |
| cART-12 | 52 | Inguinal | M | 708 | ND | 5 | EVG/COBI/3TC/TAF | 6.5 | 0 |

**Extended Data Table 2**

| **Sample ID**  **LNs** | **Imaging/**  **Panel1** | **Imaging/**  **Panel2** | **Imaging/**  **Panel3** | **Imaging/**  **Panel4** | **GeoMx** | **Visium HD** | **scRNA** | **Flow Cytometry** | **DNAscope** |
| --- | --- | --- | --- | --- | --- | --- | --- | --- | --- |
| HIVneg1 |  |  |  |  |  |  |  |  |  |
| HIVneg2 |  |  |  |  |  |  |  |  |  |
| HIVneg3 |  |  |  |  |  |  |  |  |  |
| HIVneg4 |  |  |  |  |  |  |  |  |  |
| Vir-1 |  |  |  |  |  |  |  |  |  |
| Vir-2 |  |  |  |  |  |  |  |  |  |
| Vir-3 |  |  |  |  |  |  |  |  |  |
| Vir-4 |  |  |  |  |  |  |  |  |  |
| Vir-5 |  |  |  |  |  |  |  |  |  |
| Vir-6 |  |  |  |  |  |  |  |  |  |
| Vir-7 |  |  |  |  |  |  |  |  |  |
| Vir-8 |  |  |  |  |  |  |  |  |  |
| Vir-9 |  |  |  |  |  |  |  |  |  |
| Vir-10 |  |  |  |  |  |  |  |  |  |
| Vir-11 |  |  |  |  |  |  |  |  |  |
| Vir-12 |  |  |  |  |  |  |  |  |  |
| cART-1 |  |  |  |  |  |  |  |  |  |
| cART-2 |  |  |  |  |  |  |  |  |  |
| cART-3 |  |  |  |  |  |  |  |  |  |
| cART-4 |  |  |  |  |  |  |  |  |  |
| cART-5 |  |  |  |  |  |  |  |  |  |
| cART-6 |  |  |  |  |  |  |  |  |  |
| cART-7 |  |  |  |  |  |  |  |  |  |
| cART-8 |  |  |  |  |  |  |  |  |  |
| cART-9 |  |  |  |  |  |  |  |  |  |
| cART-10 |  |  |  |  |  |  |  |  |  |
| cART-11 |  |  |  |  |  |  |  |  |  |
| cART-12 |  |  |  |  |  |  |  |  |  |

| **Epitope** | **Ab Clone** | **Fluorophore** | **Conjugated/Unconjugated** | **Cat No** | **Panel** | **Cycle** |
| --- | --- | --- | --- | --- | --- | --- |
| CD3 | OTI3E10 | Alexa-546, Opal-480 | Unconjugated | TA506064 | 1,2 | 1,2 |
| CD4 | Polyclonal | Alexa-700 | Conjugated | FAB8165N-100 | 1 | 1 |
| CD20 | L26 | eF615 | Conjugated | 42-0202-82 | 1 | 1 |
| CD8 | C8/144B | Alexa-568 | Unconjugated | M710301-2 | 1 | 1 |
| Ki67 | B56 | V450 | Conjugated | 561281 | 1 | 1 |
| PD-1 | Polyclonal | Alexa-488 | Conjugated | FAB7115G | 1 | 1 |
| CD57 | QA17A04 | BV421 | Conjugated | 393326 | 1 | 1,2 |
| TIGIT | BLR047F | Opal-620 | Unconjugated | AB243903 | 1 | 2 |
| TCF-1 | C63D9 | Opal-570 | Unconjugated | 2203S | 1 | 2 |
| ICOS | SP98 | Opal-480 | Unconjugated | AB105227 | 1 | 2 |
| GrzB | GrB-7 | Opal-650 | Unconjugated | MON7029C | 1 | 2 |
| BCL6 | GI191E/A8 | Opal-690 | Unconjugated | 760-4241 | 1 | 2 |
| CXCR-3 | 1C6 | Opal-780 | Unconjugated | 557183 | 1 | 2 |
| CD20 | L26 | Alexa546 Opal-520-480, | Unconjugated | NCL-L-CD20-L26 | 2 ,3, 4 | 1,2 |
| CD31 | WM59 | Alexa-594 | Conjugated | 303126 | 3 | 1 |
| IL-21 | Polyclonal | BV421 | Unconjugated | AHP1845 | 3 | 1 |
| FDC | CNA.42 | Alexa-488 | Unconjugated | 14-9968-82 | 3 | 1 |
| CD68 | KP-1 | Alexa-790 | Conjugated | sc-20060 AF790 | 3 | 1 |
| CD163 | EDHu-1 | Alexa-647 | Conjugated | NB110-40686AF64 | 3 | 1 |
| CD11c | EP1347Y | Alexa-555 | Conjugated | ab279329 | 3 | 1 |
| MPO | Polyclonal | Opal-690 | Unconjugated | AB9535 | 3 | 2 |
| CXCL-13 | Polyclonal | Opal-780 | Unconjugated | PA528827 | 3 | 2 |
| FOXP3 | SP97 | Alexa-594 | Conjugated | AB275080 | 2 | 1 |
| Glut1 | E4S6I | BV421 | Unconjugated | 73015 | 2 | 1 |
| CD25 | 4C9 | Alexa-488 | Unconjugated | BSB6321 | 2 |  |
| EP-2 | Polyclonal | Opal-520 | Unconjugated | PA5-33513 | 4 | 1 |
| EP-4 | C-4 | Opal-690 | Unconjugated | sc-55596 | 4 | 1 |
| COX-2 | 29 | Opal-650 | Unconjugated | sc-19999 | 4 | 1 |
| mPGES1 | Polyclonal | Opal-620 | Unconjugated | NBP2-99619 | 4 | 1 |
| Mouse IgG2b | Polyclonal | Alexa-546 | Conjugated | A-21143 | 1 | 1 |
| Mouse IgM | Polyclonal | Alexa-488 | Conjugated | 115-547-020 | 3 | 1 |
| Rabbit IgG | Polyclonal | BV421 | Conjugated | 565014 | 2,3 | 1 |
| Mouse IgG2a | Polyclonal | Alexa-546 | Conjugated | A21133 | 2 | 1 |
| Mouse IgG2b | Polyclonal | Alexa-488 | Conjugated | A21141 | 2 | 1 |

**Extended Data Table 3**

**Extended Data table 4**

| **Epitope** | **Ab Clone** | **Fluorophore** | **Cat No** | **Panel** |
| --- | --- | --- | --- | --- |
| PD1 | NAT105 | BV421 | 367422 | In vitro/PGE2-T cells |
| GITR | 108-17 | Alexa-488 | 371209 | In vitro/PGE2-T cells |
| CD4 | RPA-T4 | Alexa-700 | 557922 | In vitro/PGE2-T cells |
| ICOS | C398.4A | Alexa-647 | 313515 | In vitro/PGE2-T cells |
| CD57 | QA17A04 | PE | 393307 | In vitro/PGE2-T cells |
| CD8 | SK1 | APC/Fire750 | 344746 | In vitro/PGE2-T cells |
| Viability LIVE/DEAD™ Fixable Aqua Dead Cell Stain |  |  | L34957 | In vitro/PGE2-T cells,  Ex vivo/PLWH full panel |
| CD3 | UCHT1 | BUV805 | 612895 | Ex vivo/PLWH full panel |
| Ki67 | B56 | BUV737 | 567130 | Ex vivo/PLWH full panel |
| CXCR5 | RF8B2 | BUV615 | 751293 | Ex vivo/PLWH full panel |
| CD20 | 2H7 | BUV496 | 569672 | Ex vivo/PLWH full panel |
| CD57 | NK-1 | BUV395 | 567621 | Ex vivo/PLWH full panel |
| PD1 | EH12.2H7 | BV786 | 329930 | Ex vivo/PLWH full panel |
| CD45RO | UCHL1 | BV750 | 746942 | Ex vivo/PLWH full panel |
| CD4 | SK3 | BV650 | 563875 | Ex vivo/PLWH full panel |
| CD19 | HIB19 | BV570 | 644298 | Ex vivo/PLWH full panel |
| FOXP3 | 236A/E7 | BB700 | 566526 | Ex vivo/PLWH full panel |
| BCL6 | K112-91 | PE-Cy7 | 563582 | Ex vivo/PLWH full panel |
| GRANZYME B | GB11 | RY703 | 571462 | Ex vivo/PLWH full panel |
| CD8 | RPA-T8 | PE-CF594 | 562282 | Ex vivo/PLWH full panel |
| CD25 | M-A251 | APC-Cy7 | 561782 | Ex vivo/PLWH full panel |

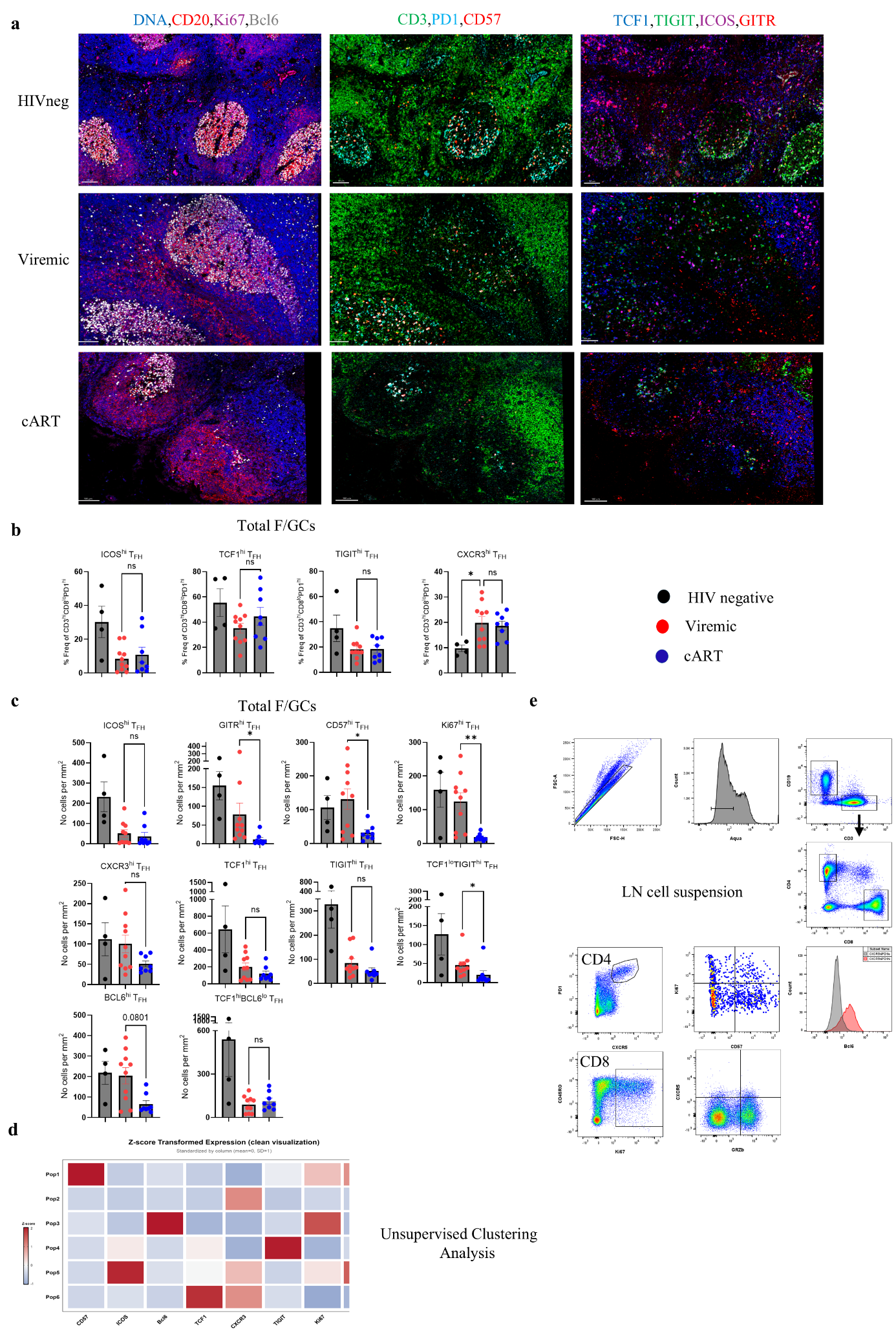
**Extended Data Figure 1.**

**a,** Representative mIF images depicting DNA/nuclei (blue), CD20 (red), Ki67 (magenta), BCL6 (grey), CD57 (red), CD3 (green), PD1(cyan), TIGIT (green), TCF1 (blue), GITR (red) and ICOS (magenta) staining in LNs from control HIV negative individuals, viremic (chronic) and cART PLWH (scale bar: 100 μm). **b,** Bar graphs showing the cell frequencies (% of CD3^hi^CD8^lo^PD1^hi^) of ICOS^hi^, TCF1^hi^, TIGIT^hi^ and CXCR3^hi^ in LN F/GCs from control HIVneg- (black, N=4), Vir (red, N=10) and cART donors (blue, N=8) quantified using imaging data via Histocytometry. Each dot represents a different donor, and bar plots show the mean ± SEM expression. **P* < 0.05(Mann-Whitney test). **c,** Bar graphs showing the cell densities (normalized per mm^2^) of CD57^hi^, TIGIT^hi^, Ki67^hi^, GITR^hi^, GITR^hi^ICOS^hi^, ICOS^hi^, TCF1^hi^, CXCR3^hi^, BCL6^hi^ and TCF1^hi^BCL6^lo^ T_FH_ in LN F/GCs from control HIVneg (black, N=4), Vir (red, N=10) and cART (blue, N=8) donors quantified using imaging data via Histocytometry. Each dot represents a different donor, and bar plots show the mean ± SEM expression. *P < 0.05 (Kruskal-Wallis ANOVA test, post-hoc Dunn΄s). **d,** Heatmap demonstrating the heterogeneity of FlowSOM identified T_FH_ cell clusters (P1-P6, generated via Cluster explorer, Flowjo v10.9.0) using the concatenated total number of cells (n=7600, 400 cells/tissue) **e,** The flow cytometry gating scheme for the identification CD3^hi^, CD19^hi^, CD4^hi^, CD8^hi^, CD4^hi^CXCR5^hi^PD1^hi^ cells and their functional phenotyping from a representative LN cell suspension is shown.

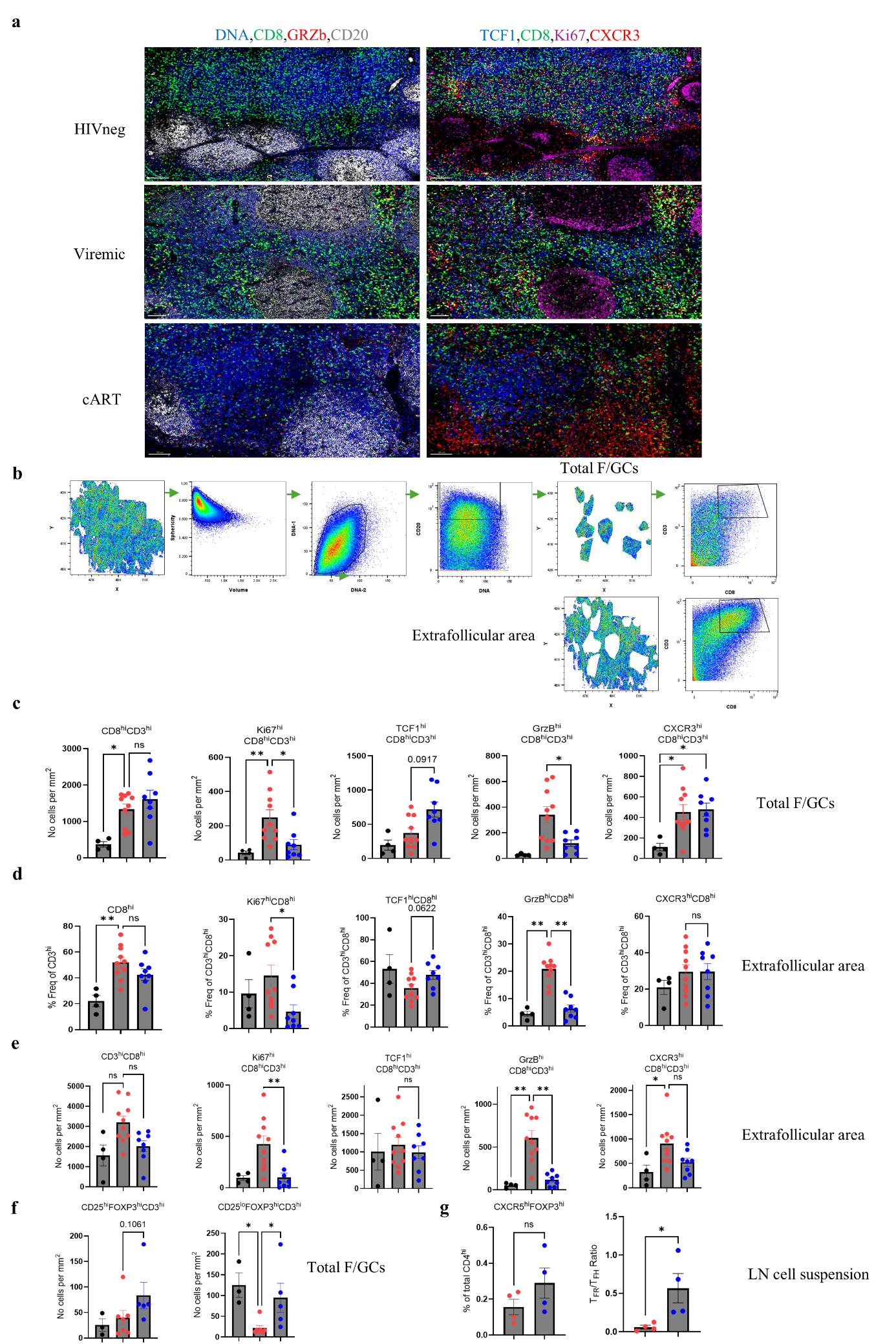

**Extended Data Figure 2.**

**a,** Representative mIF images depicting CD20 (grey), CD8 (green), GrzB (red), DNA/nuclei (blue), Ki67 (magenta), TCF1 (blue) and CXCR3 (red) staining in LNs from control HIV negative individuals, viremic (chronic) and cART PLWH (scale bar: 100 μm). **b,** The Histocytometry gating scheme for the identification of B (CD20^dim/hi^) and CD3^hi^CD8^hi^ cells in a LN is shown. Individual F/GCs were identified based on the density of CD20hi/dim cells. All follicular areas were combined (boolean gating using FlowJo10 plugin) for downstream analysis. Extrafollicular area was identified by subtracting the follicular areas from the whole tissue (boolean gating using FlowJo10 plugin). **c,** Bar graphs showing the cell densities (normalized per mm^2^) of Ki67^hi^, GrzB^hi^, TCF1^hi^ and CXCR3^hi^ CD8 cells in LN F/GCs from control HIV negative- (black, N=4), viremic (red, N=10), cART (blue, N=8) donors quantified using imaging data via Histocytometry. Each dot represents a different donor, and bar plots show the mean ± SEM expression. *P < 0.05 and **P < 0.01 (Kruskal-Wallis ANOVA test, post-hoc Dunn΄s). **d,** Bar graphs showing the cell frequencies (% of CD3^hi^ and CD3^hi^CD8^hi^) of CD8^hi^, Ki67^hi^, GrzB^hi^, TCF1^hi^ and CXCR3^hi^ in extrafollicular areas from control HIV negative- (black, N=4), viremic (red, N=10), cART (blue, N=8) donors quantified using imaging data via Histocytometry. Each dot represents a different donor, and bar plots show the mean ± SEM expression. *P < 0.05 and **P < 0.01 (Kruskal-Wallis ANOVA test, post-hoc Dunn΄s). **e,** Bar graphs showing the cell densities (normalized per mm^2^) of Ki67^hi^, GrzB^hi^, TCF1^hi^ and CXCR3^hi^ CD8 cells in extrafollicular areas from control HIV negative- (black, N=4), viremic (red, N=10) and cART (blue, N=8) donors quantified using imaging data via Histocytometry. Each dot represents a different donor, and bar plots show the mean ± SEM expression. *P < 0.05 and **P < 0.01 (Kruskal-Wallis ANOVA test, post-hoc Dunn΄s). **f,** Bar graphs showing the cell densities (normalized per mm^2^) of CD25^hi^Foxp3^hi^ and CD25^lo^Foxp3^hi^ in LN F/GCs from control HIV negative- (black, N=3), viremic (red, N=7) and cART (blue, N=5) donors quantified using imaging data via Histocytometry. Each dot represents a different donor, and bar plots show the mean ± SEM expression. **P* < 0.05 (Kruskal-Wallis ANOVA test, post-hoc Dunn΄s). **g,** Bar graphs demonstrating the cell frequency (% of CD4^hi^) of CXCR5^hi^Foxp3^hi^ and the T_FR_ (CXCR5^hi^Foxp3^hi^) /T_FH_ (CXCR5^hi^PD^hi^) ratio in Vir (red, N=4) and cART (blue, N=4) LN cell suspensions samples as measured by Flow Cytometry. Each dot represents a different donor, and bar plots show the mean ± SEM expression. **P* < 0.05 (Mann-Whitney test).

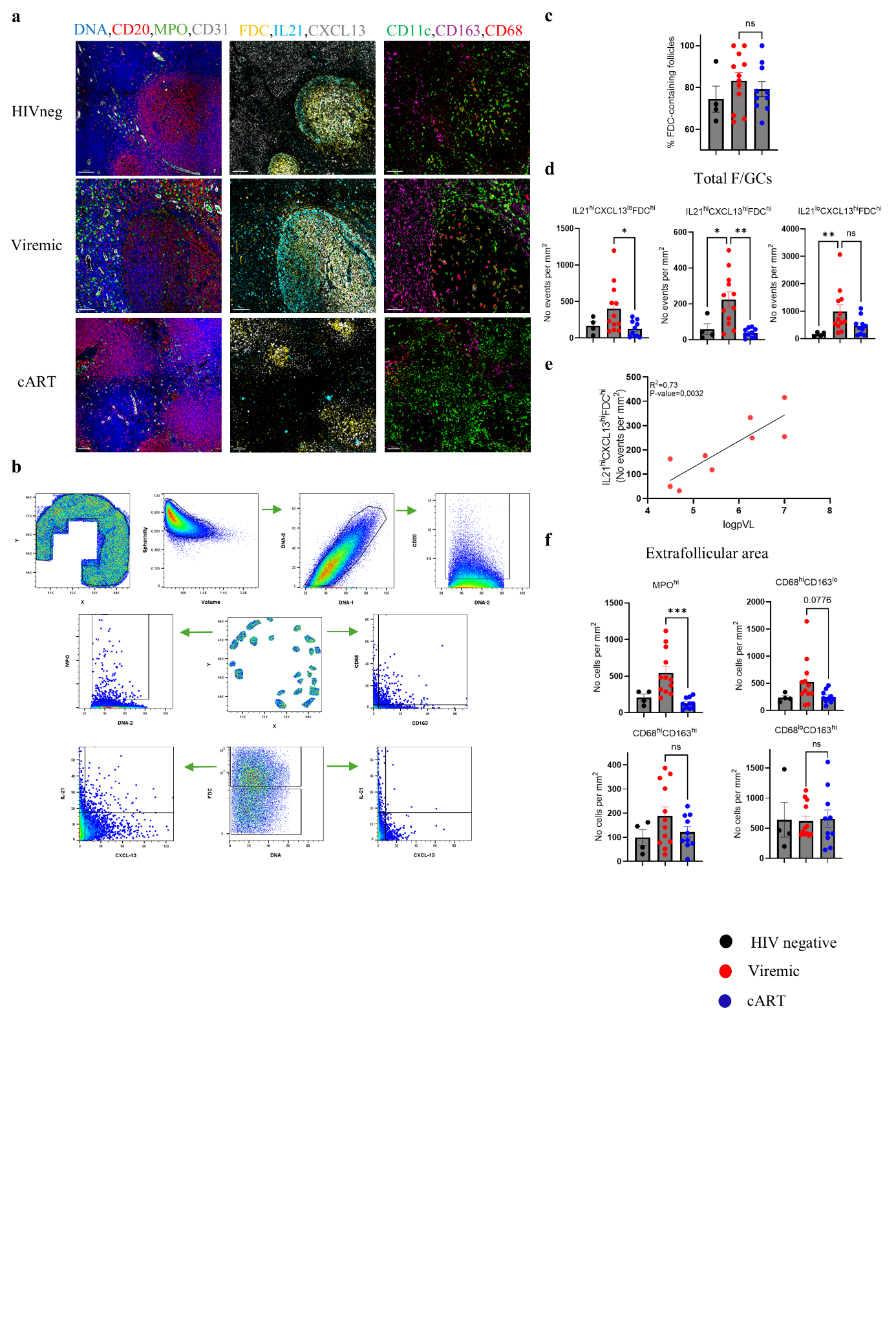

**Extended Data Figure 3.**

**a,** Representative mIF images depicting CD20 (red), MPO (green), CD31 (grey), DNA/nuclei (blue), FDC (yellow), IL21 (cyan), CXCL13 (grey), CD163 (magenta), CD11c (green) and CD68 (red) staining in LNs from control HIV negative individuals, viremic (chronic) and cART PLWH (scale bar: 100 μm). **b,** The Histocytometry gating scheme for the identification of B (CD20^hi/dim^), IL21^hi^CXCL13^lo^FDC^lo^, IL21^hi^CXCL13^hi^FDC^lo^, IL21^lo^CXCL13^hi^FDC^lo^, MPO^hi^, CD68^hi^CD163^lo^, CD68^hi^CD163^hi^ and CD68^lo^CD163^hi^ cells in a LN is shown. Individual F/GCs were identified based on the density of CD20^hi/dim^ cells. All follicular areas were combined (boolean gating using FlowJo10 plugin) for downstream analysis. **c,** Bar graph showing the number of intact FDC-structures in different LN F/GCs from control HIVneg (black, N=4), Vir (red, N=12) and cART (blue, N=10) donors as calculated by manual inspection. Each dot represents a different donor, and bar plots show the mean ± SEM expression. ns>0,05 (Kruskal-Wallis ANOVA test, post-hoc Dunn΄s). **d,** Bar graphs showing the cell densities (normalized per mm^2^) of IL21^hi^CXCL13^lo^FDC^hi^, IL21^hi^CXCL13^hi^FDC^hi^, IL21^lo^CXCL13^hi^FDC^hi^ events in LN F/GCs from control HIVneg (black, N=4), viremic (red, N=12) and cART (blue, N=10) donors. Each dot represents a different donor, and bar plots show the mean ± SEM expression. **P* < 0.05 and **P < 0.01 (Kruskal-Wallis ANOVA test, post-hoc Dunn΄s). **e,** Linear regression analysis between IL21^hi^CXCL13^hi^FDC^hi^ events and peripheral blood viral load (logpVL). R^2^ and p-value are listed.  Each dot represents a different donor. **f,** Bar graphs showing the cell densities (normalized per mm^2^) of MPO^hi^, CD68^hi^CD163^lo^, CD68^hi^CD163^hi^, CD68^lo^CD163^hi^, CD11c^hi^ and CD11c^hi^CD68^hi^ cells in LN extrafollicular areas from control HIVneg (black, N=4), viremic (red, N=12) and cART (blue, N=10) donors. Each dot represents a different donor, and bar plots show the mean ± SEM expression. ***P < 0.001 (Kruskal-Wallis ANOVA test, post-hoc Dunn΄s).

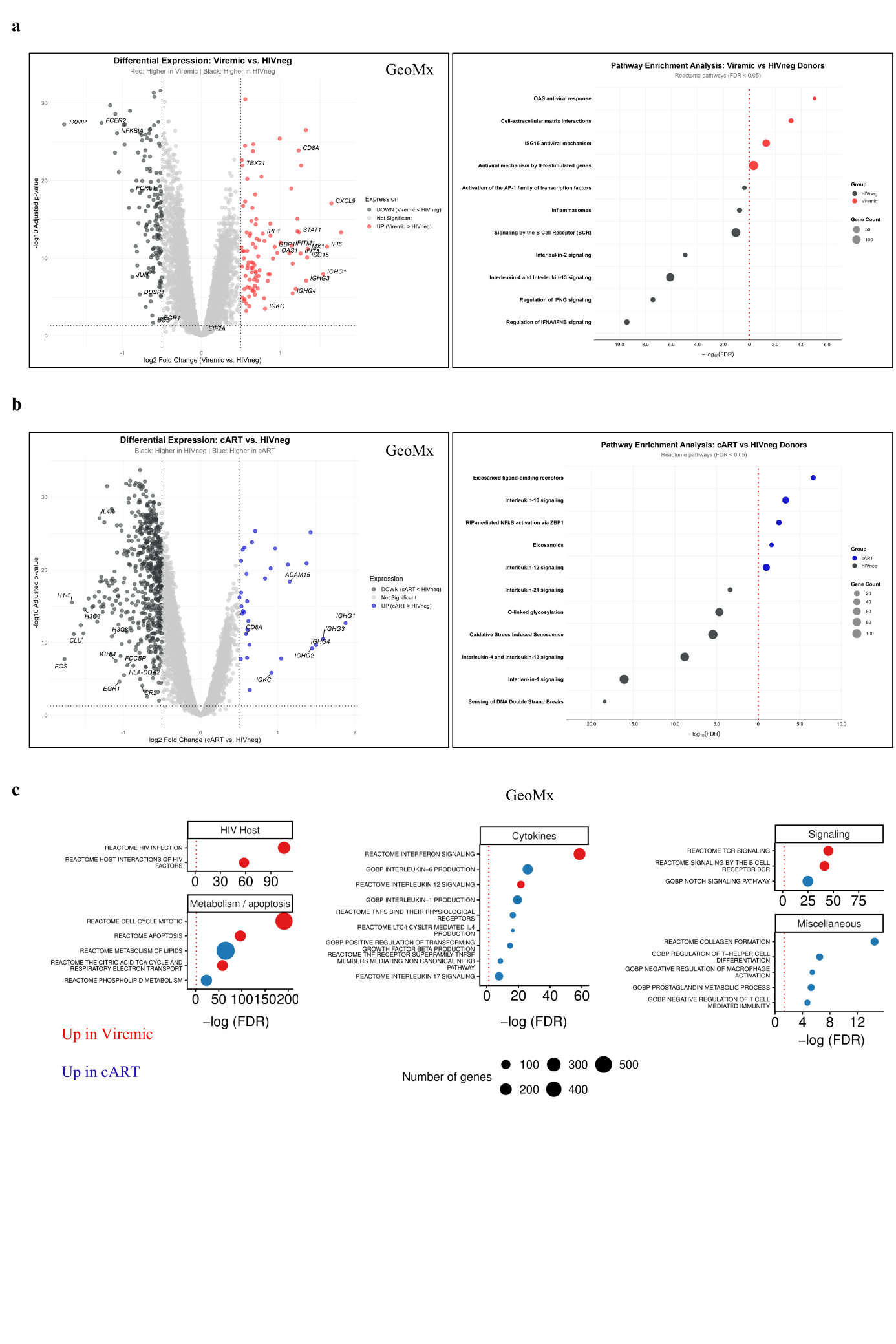

**Extended Data Figure 4.**

**a,** (left panel) Volcano plot showing differentially expressed genes between control HIVneg and Vir LN follicular ROIs. Red dots indicate genes upregulated in Vir, black dots in control HIV negative, and grey dots denote non-significant genes. Vertical ticked lines represent the fold-change threshold. (right panel) Bubble plot of gene pathways significantly enriched in Vir- (red) versus control HIVneg LN follicular ROIS (black). Bubble size indicates the number of differentially expressed genes for every pathway. The x-axis shows the negative logarithm of the FDR **b,** (left panel) Volcano plot showing differentially expressed genes between control HIVneg and cART LN follicular ROIs. Blue dots indicate genes upregulated in cART, black dots in control HIVneg, and grey dots denote non-significant genes. Vertical ticked lines represent the fold-change threshold. (right panel) Bubble plot of gene pathways significantly enriched in cART LN- (blue) versus control HIVneg LN follicular ROIs (black). Bubble size indicates the number of differentially expressed genes for every pathway. The x-axis shows the negative logarithm of the FDR. **c,** Bubble plots of gene pathways related to HIV, Metabolism/Apoptosis, Cytokines and T-B cell Signalling significantly enriched in Vir (red) versus cART F/GCs (blue). Bubble size indicates the number of differentially expressed genes for every pathway. The x-axis shows the negative logarithm of the FDR.

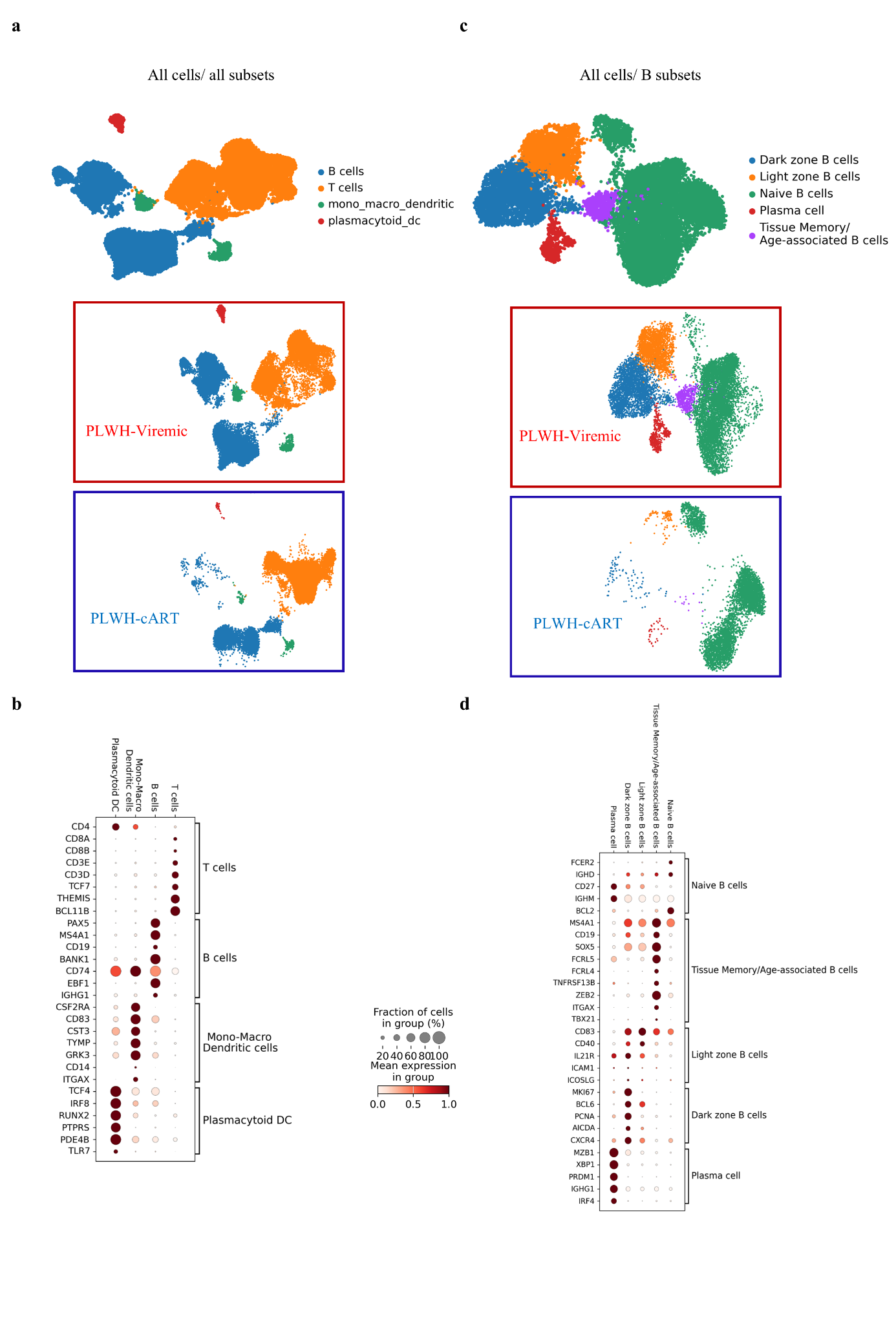

**Extended Data Figure 5.**

**a,** UMAP visualization of unsupervised clustering of total LN cells (all cells, Vir and cART, N=8, n=42936), Vir (N=4, n=27209) and cART (N=4, n=15727) as determined by scRNAseq, resulting in 6 different clusters. **b,** Dot plot demonstrating the expression of genes characteristic of different bulk cell subsets. The colour of the dot indicates the mean expression intensity, while the size of the sphere shows the percentage of cells expressing each gene. **c,** UMAP visualization of unsupervised clustering of total LN B cells (all cells, Vir and cART, N=8, n=17622), Vir (N=4, n=12975) and cART (N=4, n=4647) as determined by scRNAseq, resulting in 6 different clusters. **d,** Dot plot demonstrating the expression of genes characteristic of different B cell subsets. The colour of the dot indicates the mean expression intensity, while the size of the sphere shows the percentage of cells expressing each gene.

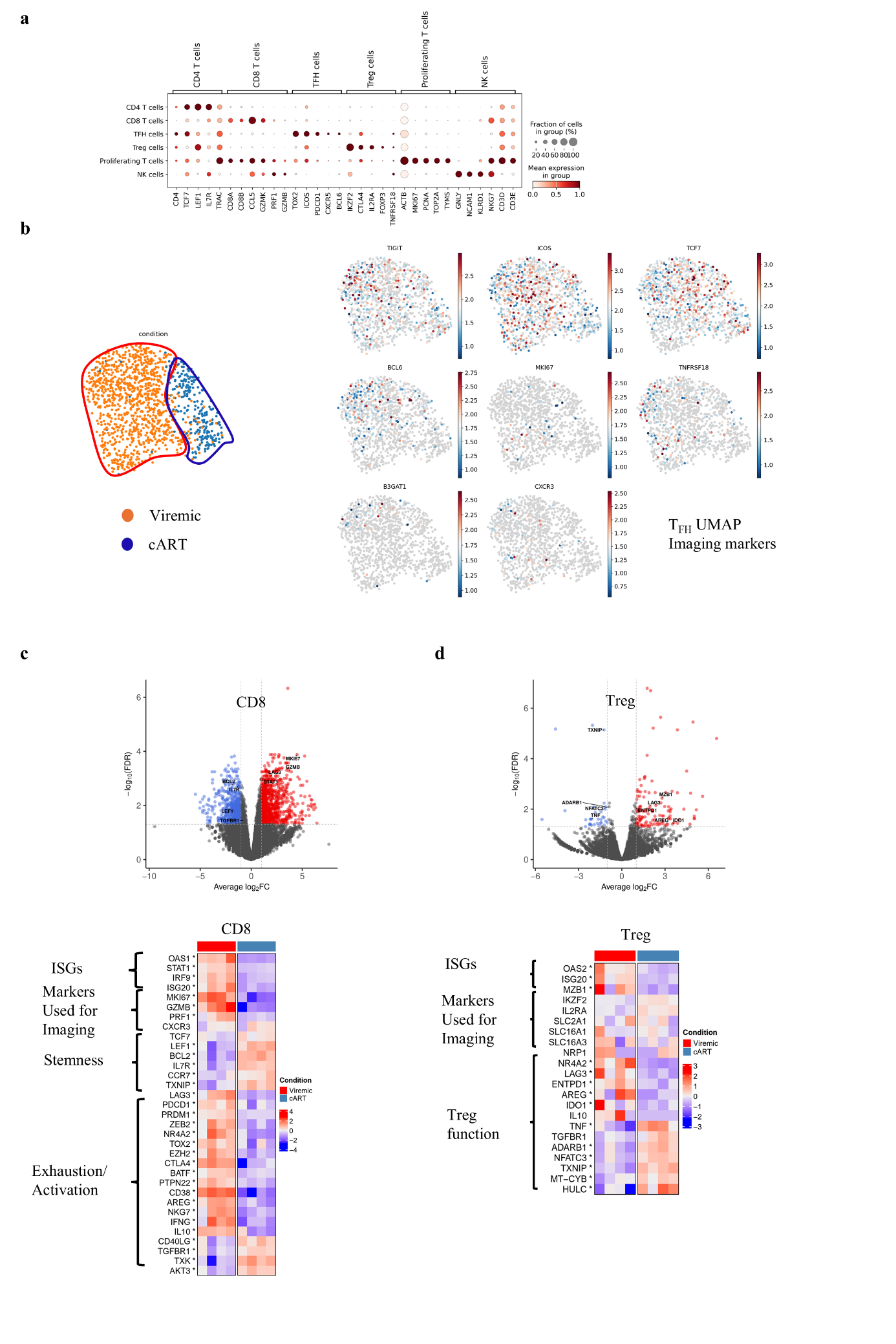

**Extended Data Figure 6.**

**a,** Dot plot showing the expression of genes characteristic of different T cell subsets. The colour of the dot indicates the mean expression intensity, while the size of the sphere shows the percentage of cells expressing each gene. **b,** UMAP displaying unsupervised clustering of total T_FH_ cells (all cells, Vir and cART, N=8, n=1212), Vir (N=4, n=972) and cART (N=4, n=240) as determined by scRNAseq. The mRNA expression levels of the protein markers used for mIF analysis are also shown (log-normalized and clipped at the 99th percentile). **c-d,** Volcano plot of differentially expressed genes between Vir and cART LN CD8 **(c)** and T_regs_ (**d)** cells (upper panel). Vertical ticked lines indicate the fold-change threshold. Heatmaps displaying differentially expressed genes among CD8 **(c)** and T_regs_ **(d)** (lower panel) cells. Genes displayed on the y-axis are classified into functional categories as noted. Upregulated genes are displayed in red and downregulated genes in blue based on their z-score.

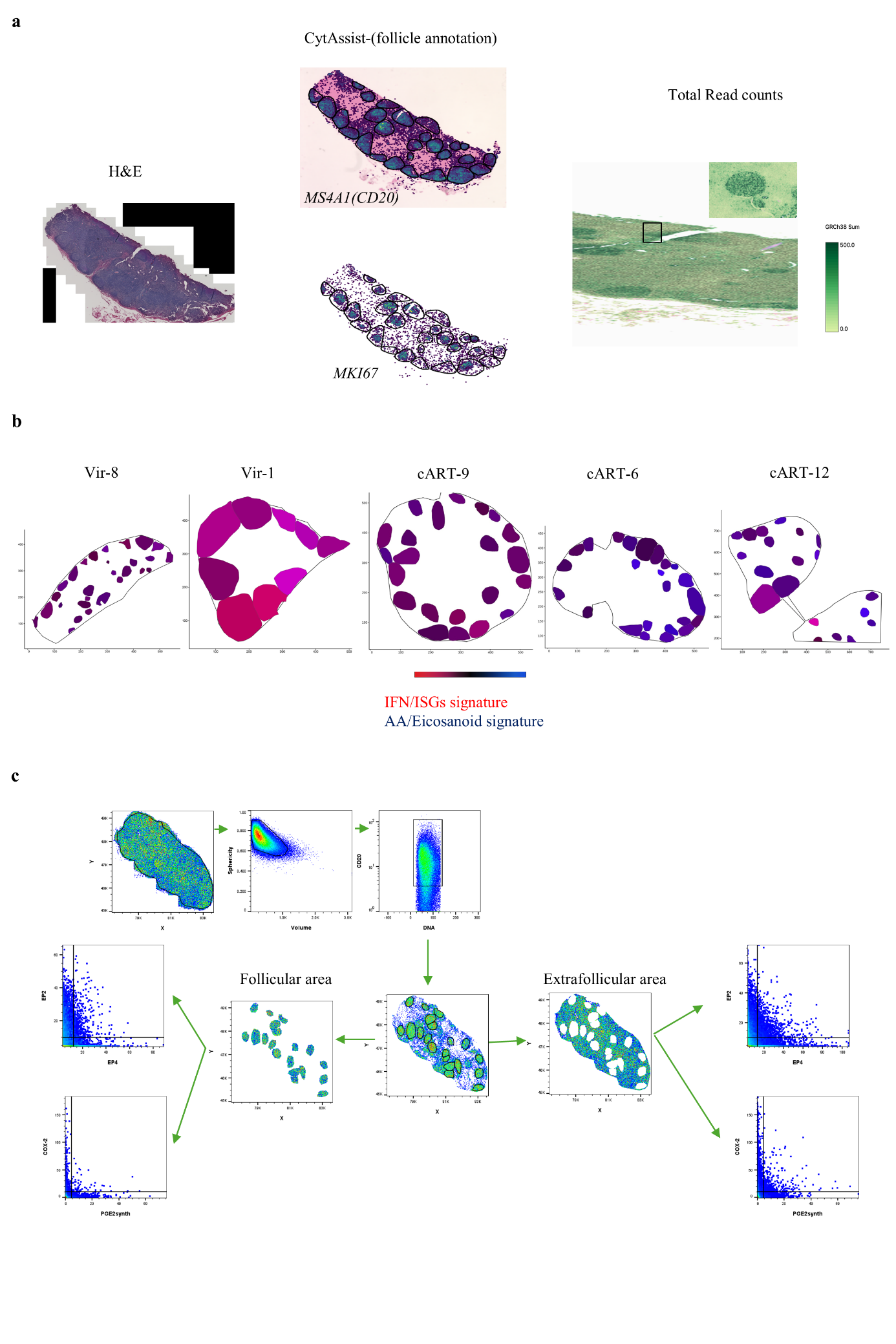

**Extended Data Figure 7.**

**a,** Cartoon displaying representative images of an analysed tissue (H&E, CytAssist, Loope Browser). ROIs (follicular areas) were manually annotated using microscopy images and CD20/MS4A1 mRNA expression. Simultaneous MKI67 in-situ expression was indicative of secondary follicles. Segmentation of LN cells was conducted using the H&E image by applying SpaceRanger. **b,** Representative 2D tissue plots illustrating the spatial gradient of gene signature enrichment across F/GCs from viremic and cART donors, as calculated by UCell. Red indicates enrichment of the IFN/ISG signature, whereas blue indicates enrichment of the AA/Eicosanoid signature. See legend for Fig 6B for further details. **c,** The Histocytometry gating scheme for the identification of EP2^hi^EP4^lo^, EP2^hi^EP4^hi^, EP2^lo^EP4^hi^, COX2^hi^ and PGES1^hi^ cells in a representative LN is shown. Individual F/GCs were identified based on the density of CD20^hi/dim^ cells. All follicular areas were combined (boolean gating using FlowJo10 module) for downstream analysis. Extrafollicular space was identified by excluding the follicular regions using the boolean module.

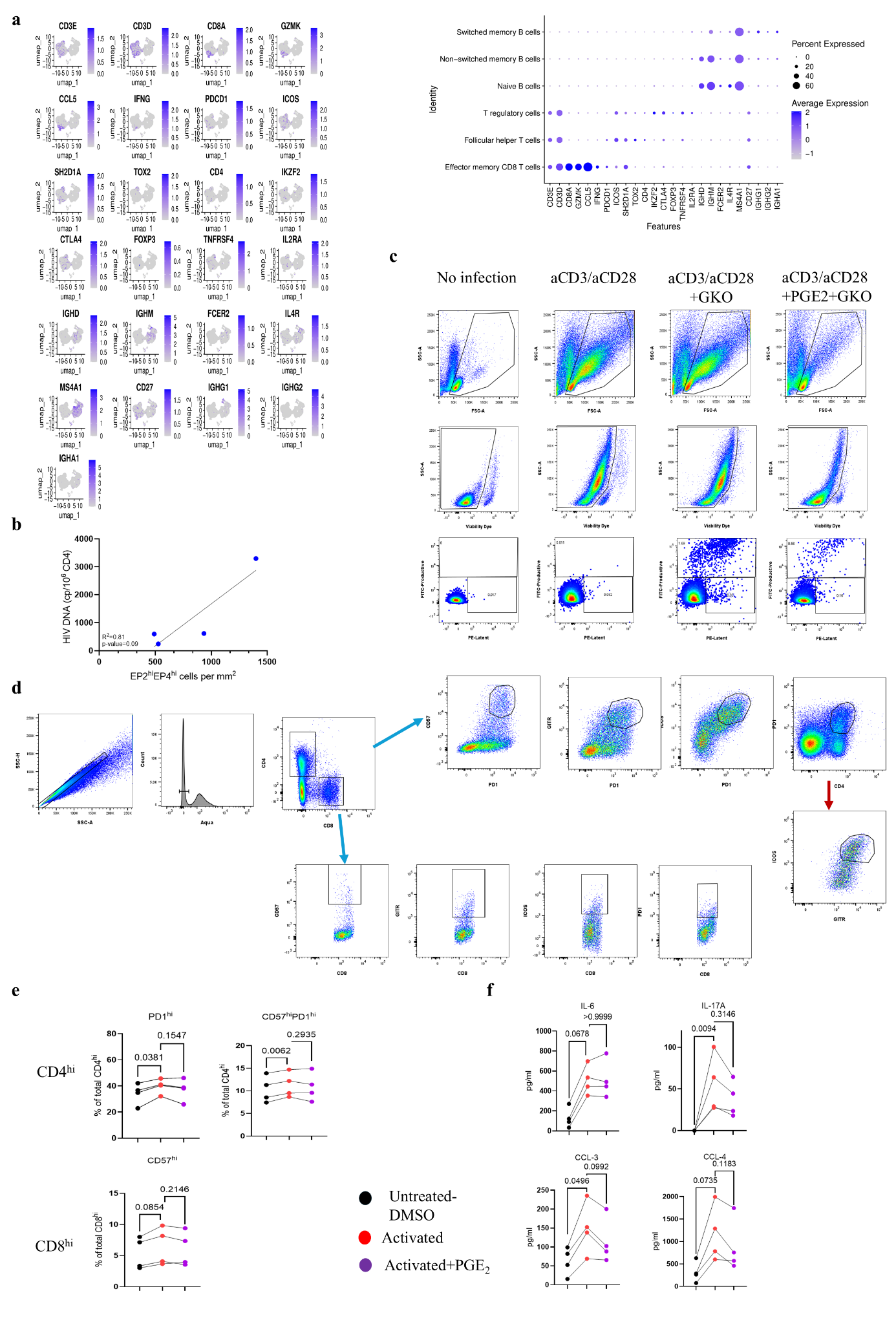
**Extended Data Figure 8.**

**a,** Dot plot showing the expression of genes characteristic of different analysed cell subsets from an independent viremic PLWH cohort (Neutralizers, Non-Neutralizers) as determined by scRNAseq (right). The colour of the dot indicates the mean expression intensity, while the size of the sphere shows the percentage of cells expressing each gene. UMAP feature plots demonstrating the expression patterns of selected lineage- and activation-associated genes (left). **b,** Linear regression analysis between EP2^hi^EP4^hi^ cell densities of cART (blue, N=4) F/GCs and matched PBMCs HIV DNA measurements (cp/10^6^ CD4 cells). R^2^ and p-values are listed. Each dot represents a different donor. **c,** Flow cytometry gating scheme for the identification of *in vitro* Productively (FITC) - and Latently (PE)- GKO infected cells upon activation with or without PGE_2_ administration. Non-infected control is also shown. **d,** Representative flow cytometry gating strategy for the identification of activated CD4^hi^ and CD8^hi^ T-cell subsets based on the expression of PD-1, ICOS, GITR, and CD57 in a primary tonsillar cell suspension. **e,** Graphs demonstrating the cell frequency (%) of control (DMSO) and activated, with or without concomitant PGE_2_ administration, CD4 and CD8 subsets based on PD1 and CD57expression (N=4). P-values are also shown. **P* < 0.05 ***P < 0.001 (one-way ANOVA tests) **f,** Graphs demonstrating the concentration of IL-8, IL-17A, CCL-3 and CCL-4 in supernatants from control (DMSO) and activated, with or without concomitant PGE_2_ administration, human primary tonsillar cells (N=4). P-values are also shown. **P* < 0.05 (one-way ANOVA tests)

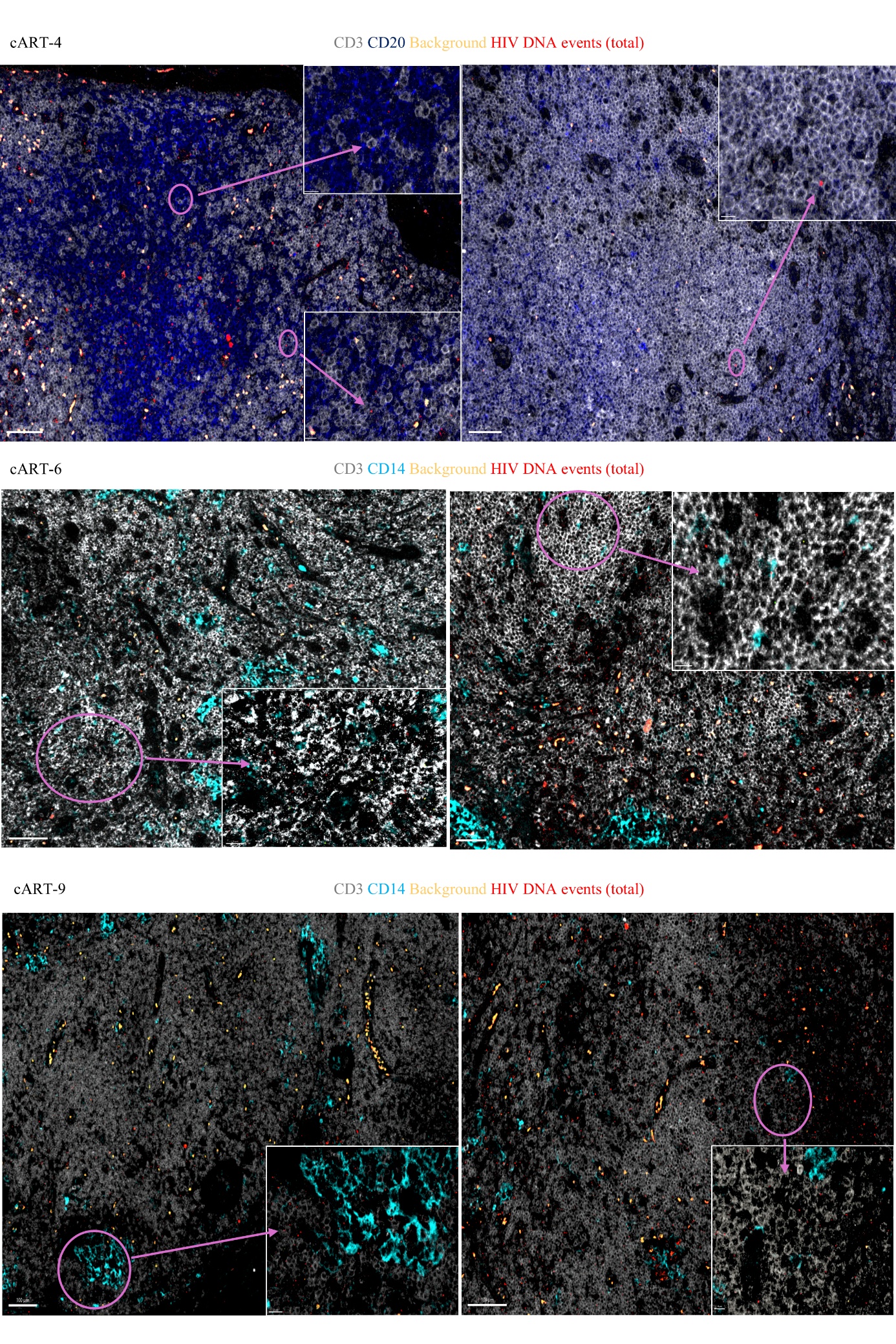
**Extended Data Figure 9.**

Representative mIF images showing CD20 (blue), CD3 (grey), CD14 (cyan), background events (dark yellow) and HIV DNA events (red) in three different LNs from cART PLWH (scale bar: 70-100 μm). Zoomed areas are shown as insertions.

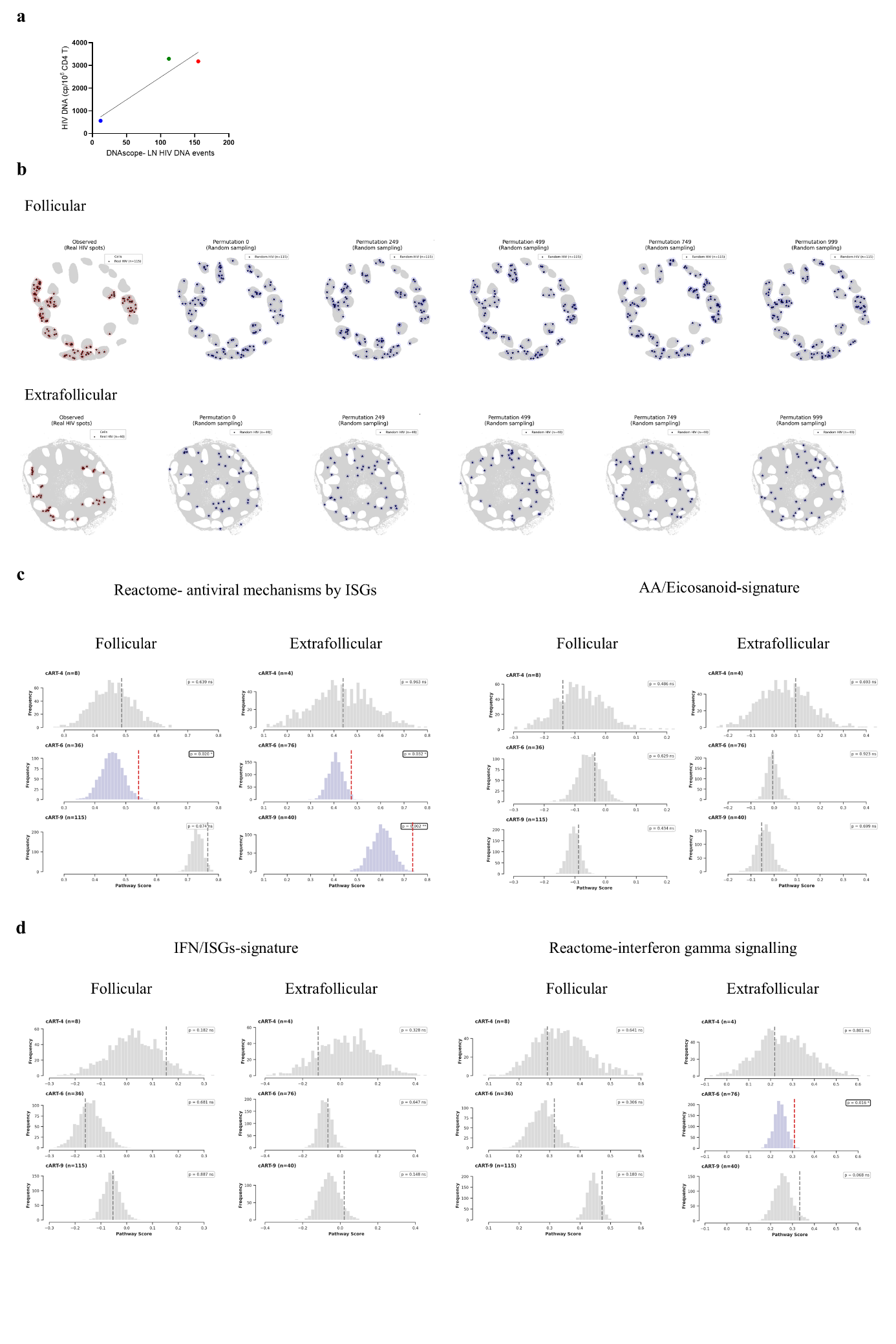
**Extended Data Figure 10.**

**a,** Linear regression analysis between the total number of HIV DNAscope-detected latently infected CD3^hi^ cells from cART (N=3) LNs and matched PBMCs HIV DNA measurements (cp/10^6^ CD4 cells). Different donors are colour coded. **b,** Representative example of permutation-based spatial neighbourhood analysis of follicular (upper panel) and extrafollicular (lower panel) DNAscope-generated HIV DNA+ events (red stars). Randomly positioned (n=1000) HIV DNA events (blue stars) are also shown. **c,** Histogram showing distributions of random and observed follicular and extrafollicular pathway scores (Reactome-Antiviral mechanisms by ISGs[left], in-house curated AA/Eicos signature[right]) in LNs from cART PLWH. Dashed red lines indicate the observed pathway score, and p values were computed for the observed score relative to the distribution of random permutation scores. Each donor’s HIV DNA events count (n) is also listed. **d,** Histogram distributions of follicular and extrafollicular pathway scores (in-house curated IFN/ISG gene signature[left], Reactome-Interferon gamma signalling [right]) in LNs from cART PLWH. Dashed red lines indicate the observed pathway score, and p values denote comparisons with the reference distribution. Each donor’s HIV DNA events count (n) is also listed.
